# Understanding the genetic diversity of the guayabillo (*Psidium galapageium*), an endemic plant of the Galapagos Islands

**DOI:** 10.1101/2020.09.28.317602

**Authors:** Diego Urquia, Gabriela Pozo, Bernardo Gutierrez, Jennifer K. Rowntree, Maria de Lourdes Torres

## Abstract

Oceanic archipelagos are known to host a variety of endemic plant species. The genetic diversity and structure of these species is an important indicator of their evolutionary history and can inform appropriate conservation strategies that mitigate the risks to which they’re exposed, including invasive species and environmental disturbances. A comprehensive consideration of the role of their natural history, as well as the landscape features and the geological history of the islands themselves is required to adequately understand any emerging patterns. Such is the case for the guayabillo (*Psidium galapageium*), an understudied endemic plant from the Galapagos Islands with important ecological and economic roles. In this study we designed and evaluated 13 informative SSR markers and used them to investigate the genetic diversity, population structure and connectivity of the guayabillo populations from San Cristobal, Isabela and Santa Cruz islands. A total of 208 guayabillo individuals were analyzed, revealing a strong population structure between islands and two distinct genetic lineages for the Santa Cruz population. Overall, the guayabillo genetic diversity is relatively high, an unusual pattern for an insular endemic species which is possibly explained by its polyploidy and the geographical features of the islands. These include their broad altitudinal ranges and habitat heterogeneity. For populations displaying a lower genetic diversity such as San Cristobal, the history of human disturbance could be an important factor explaining these observations. Some similarities between individuals in Santa Cruz and the San Cristobal population could be explained by population differentiation or distinct natural histories of separate lineages. Our findings highlight the complex population dynamics that shape the genetic diversity of species like the guayabillo and emphasize the need to explore the currently unresolved questions about this Galapagos endemic plant.

## 1. Introduction

Oceanic islands are home to unique species which have emerged as a product of their evolutionary histories being driven by geographical isolation and distinct topological and climatic conditions. This makes them ideal study cases for evolutionary and ecological processes (Carlquist, 1974; Emerson, 2002; Shaw and Gillespie, 2016). Studying these species has been an important step in addressing evolutionary biology questions about key processes such as adaptation, speciation, radiation, and the link between evolution and geography (Geist et al., 2014; Rumeu et al., 2016; Shaw and Gillespie, 2016). Among these, insular endemics are an interesting case of species that comprise distinct gene pools compared to their counterparts in mainland ecosystems. The genetic diversity patterns observed for insular organisms are driven by factors that include founder events and genetic bottlenecks, phenomena that island species usually experience at some point in their history (Mayr, 1954; Hagenblad et al., 2015; Stuessy et al., 2014). The characteristics of the island inhabited by specific plant populations or species, including their size, age and habitat heterogeneity, are also major factors that determine their genetic diversity (MacArthur and Wilson, 1967; Stuessy et al., 2014).

The Galapagos Islands are a prime example of these oceanic archipelagos; they are conformed by 13 main islands and more than 100 minor islets of volcanic origin. The archipelago is located in the Pacific Ocean, ~1000 km off the coast of South America. Thanks to their tropical location and oceanographic situation, the Galapagos harbor a great variety of unique species, as well as rich ecosystems which remain relatively undisturbed compared to other insular systems (Gillespie and Clague, 2009; Jaramillo et al., 2011). Moreover, the overall young age of the archipelago and the coexistence of islands of different ages provide a *real-time* observation window of evolutionary processes (Jaramillo et al., 2011).

The endemic species of the Galapagos have been extensively studied in the context of their evolution and conservation. However, most research has been focused on animal species (Geist et al., 2014; Shaw and Gillespie, 2016). Few studies explore the genetic diversity and population structure of endemic plant species, both being key properties for understanding the evolutionary history and assessing the vulnerability and responsiveness to environmental change of the archipelago’s endemic flora (Fridley et al., 2007; Jump et al., 2009; Stuessy et al., 2014). In fact, insular endemic species are valuable genetic resources for scientific development in fields such as bioprospection and plant breeding (e.g. Guezennec et al., 2006; Pailles et al., 2017). Unfortunately, endemic insular species are intrinsically vulnerable to threats which include environmental change, disease, invasive species, human perturbation and habitat loss due to their isolation, relatively small population sizes and restricted distribution (Whittaker, 1998; Sakai et al., 2001). Thus, it is not surprising that in 2016, 40% of all recognized endangered species were found in island ecosystems (Island Conservation, 2016). The identification of factors that promote or disfavor genetic diversity, and the assessment of genetic structure, also assist in the establishment of conservation areas, the prioritization of populations for conservation action and the adequate evaluation of such strategies (Bensted-Smith, 2002; Wallis and Trewick, 2009; Moritz, 2002; Gitzendanner et al., 2012).

Multiple driving forces have been associated with the evolution and genetic diversity of the endemic species in the Galapagos Islands. For instance, *Scalesia affinis* presents a higher genetic diversity in Isabela island compared to Floreana island, partially explained by the former having a much larger landmass and a broader altitudinal gradient (Nielsen, 2004). Other factors pertaining to the evolutionary history of the species, including speciation mechanisms (anagenesis vs. cladogenesis) and other events such as past hybridization and polyploidization, should also be considered for interpreting genetic diversity patterns (Soltis and Soltis, 2000; Stuessy et al., 2006; Stuessy et al., 2014). It has been proposed, for example, that the Galapagos endemic shrub *Galvezia leucantha* harbors high levels of genetic diversity in part due to populations from different islands maintaining some gene flow (Guzmán et al., 2016); thus, all these populations still conform a single species (as observed in anagenesis) (Stuessy et al., 2014; Takayama et al., 2015). Furthermore, the reproductive biology (outcrossing vs. selfing vs. clonal reproduction) and dispersal mechanisms of the species are also relevant factors that explain genetic diversity and structure (Crawford and Whitney, 2010). Species that inbreed, self-pollinize and/or reproduce clonally tend to show higher levels of genetic differentiation among populations, especially if they are weak dispersers (Ellstrand and Elam, 1993; Hamrick and Godt, 1996). For instance, the low heterozygosity and high between-population differentiation in the Galapagos endemics *Solanum cheesmaniae* and *Solanum galapagense* were partially attributed to their highly autogamous nature (Rick, 1983; Pailles et al., 2017). On the other hand, it is thought that gynodioecious dimorphism in *Lycium minumum* emerged as a mechanism to promote outcrossing and to maintain genetic diversity; in turn, this dimorphism would be linked with a tetraploidization event in the evolutionary history of the species (Sakai et al. 1995; Levin et al., 2015).

More interestingly for this particular case, the recent geological history of the Galapagos Islands themselves must be considered when interpreting and understanding the genetic diversity and structure of an endemic plant species. Every island of the archipelago emerged progressively due to the eastward movement of the Nazca Plate over a mantle hotspot (Villagómez et al., 2007; Geist et al., 2014); thus, the older islands of the archipelago are located to the southeast, while the newest ones are located to the northwest (Geist et al., 2014). This movement of the Nazca Plate, in combination with historical changes in the sea level, lead to oceanic barriers that separated islands that emerged over the same hotspot and were initially close together (Christie et al., 1992; Geist et al., 2014). In consequence, populations from different islands are kept separated from each other by considerable stretches of ocean extending for several kilometers. Moreover, these isolated populations may be exposed to different environmental conditions and to different demographic events and genetic processes (e.g. population size changes, selection, genetic drift, mutations, etc.) (Lombaert et al., 2011; Shirk et al., 2014), establishing distinct patterns of genetic structure within a species and even triggering speciation (Rumeu et al., 2016; Pailles et al., 2017). This phenomenon has been observed in Galapagos endemic plants such as *S. cheesmaniae* and *L. minimum,* where a notorious genetic divergence arose between populations of the older eastern islands and the western younger islands (Levin et al., 2015; Pailles et al., 2017).

Guayabillo (*Psidium galapageium;* Myrtaceae) is one of the 241 endemic plant species in the Galapagos Islands (Jaramillo et al., 2014). Catalogued as *Near threatened* in the Red Book of endemic plants of Ecuador (Kawasaki et al., 2017), it is one of the few endemic tree-like plants in the archipelago, and hence a significant landscape component of the transition zones and *Scalesia* forests of several islands (San Cristobal, Santa Cruz, Santiago, southern Isabela, Fernandina, Pinta and Floreana); its distribution also includes drier lowland and humid highland sites (Porter, 1968; McMullen, 1999). Guayabillo serves as an anchoring substrate for nutrient-fixing lichen (Dal Forno et al., 2017), and chemical compounds produced by its leaves have been used as a natural repellent for parasitic and hematophagous insects by birds, including several species of endemic finches (Cimadom et al., 2016). Its hard and resistant wood is used by the islanders for house and boat construction (Wiggins et al., 1971). Nevertheless, as many of the endemic plants of the Galapagos, guayabillo is threatened by human-induced disturbances including overexploitation of its wood, habitat loss, and the presence of invasive species (Wiggins et al., 1971; Adsersen et al., 1988; Frankham, 1995; Tye et al., 2007; Dal Forno et al., 2017). The direct competition between endemic and invasive species can cause a reduction and fragmentation in the populations of the former, as well as a loss of its genetic diversity (Nielsen, 2004; Jaramillo et al., 2011; Stuessy et al., 2014). For this reason, the introduction of exotic species is of great concern in insular ecosystems like the Galapagos (Whittaker, 1998; Tye et al., 2007). The common guava (*Psidium guajava*), for example, is an invasive species that shares some of the same ecosystems with guayabillo, raising the potential risk of guava populations outcompeting or forming interspecific hybrids with its endemic relative (which could cause genetic erosion) (Torres and Gutiérrez, 2018). Similarly, the Galapagos flora in general is threatened by destructive grazers such as goats and feral livestock; these animals have already caused an impact for several endemic species in the islands such as *Calandrina galapagosa, S. affinis* and *G. leucantha* (Nielsen, 2004; Jaramillo et al., 2011; Guzmán et al., 2016). Despite its economic and ecological importance and potential vulnerability as an island endemic, little is known about the natural history of the guayabillo and its population genetics.

We present the design and evaluation of homologous SSR primers for *P. galapageium* in order to assess the genetic diversity, structure and connectivity of three populations of this species, in San Cristobal, Isabela and Santa Cruz Islands. The parameters inferred from the genetic data are used to describe the natural history of the species in the archipelago in combination with the currently available data regarding the biology of guayabillo, the geography of the Galapagos and their geological history, and the presence of human populations in the islands. This knowledge can be used to identify potential risks to guayabillo populations in the archipelago, and is relevant for the establishment and evaluation of conservation strategies.

## 2. Material and methods

### 2.1. Study sites and sample collection

In order to identify *P. galapageium* individuals, the morphological description by Porter (1968) was used. Guayabillo is a small tree or shrub of smooth, pinkish gray bark (Fig. 1a). Its branches are divaricate, its branchlets terete and gray. Its leaves are elliptic to ovate, equilateral and 1.8-5.5 cm long and 0.9-2.6 cm wide. Flowers are 1-1.5 cm in diameter, of a whitish color (Fig. 1b). Berries have a 2 cm diameter, they are globose to subglobose, glabrous, and of a pale yellow to yellow color (Fig. 1c).

**Fig. 1.**
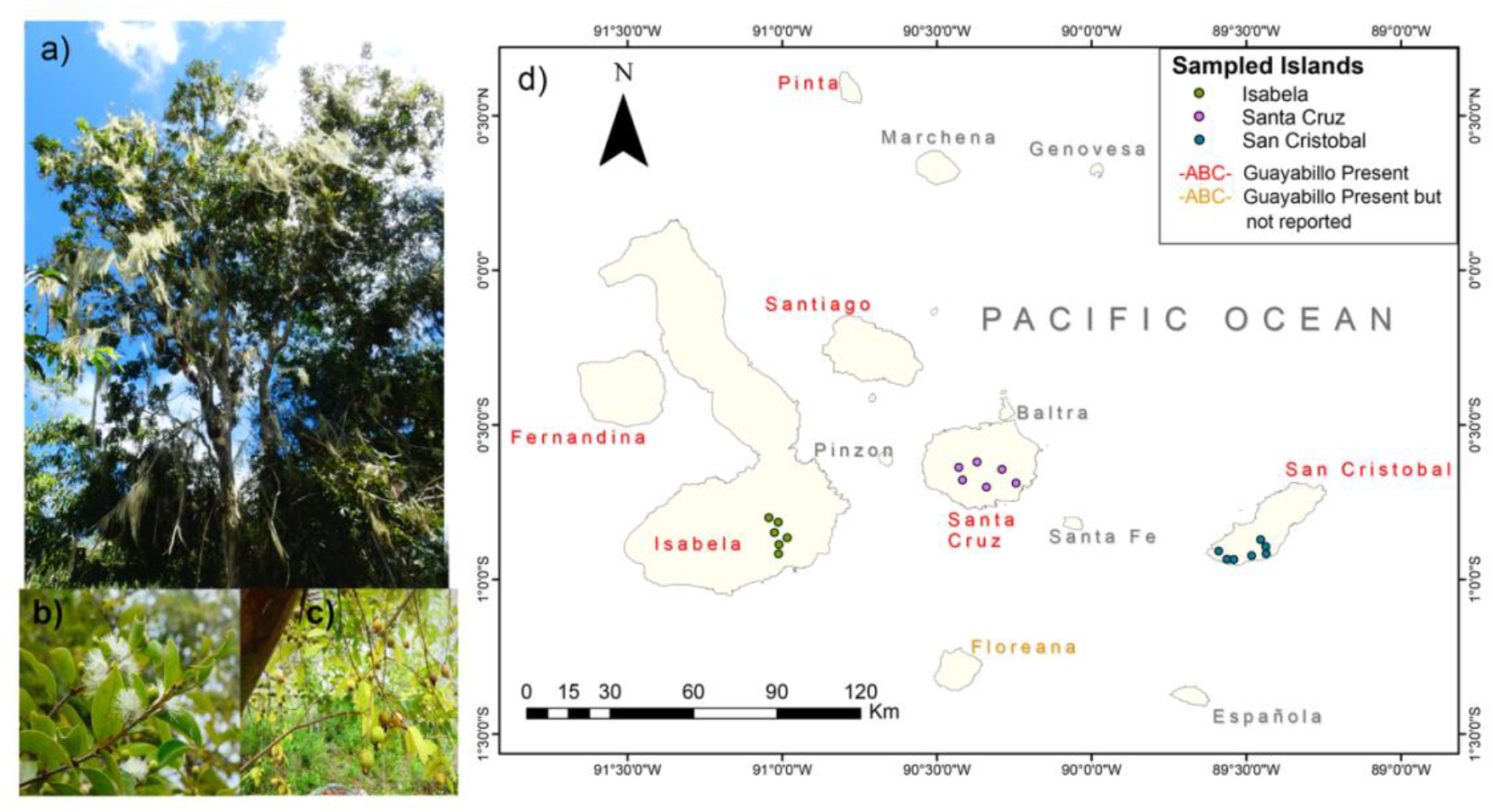
a) A guayabillo tree, b) Details of the leaves and flowers of guayabillo. c) Details of the leaves and fruits of guayabillo. (Photos: Bryan Reatini, UNC-CH). d) Galapagos Islands map indicating the sampling sites of this study in Isabela, Santa Cruz and San Cristobal Islands. The islands where guayabillo is distributed are highlighted in red; note that although guayabillo is not officially reported as present in Floreana Island (orange label), it is actually distributed over there as well (Bryan Reatini, pers. comm.)

Samples from *P. galapageium* individuals were collected from three islands: San Cristobal (seven sampling locations), Santa Cruz (six sampling locations) and Isabela (six sampling locations; Fig. 1d). For the selection of these sampling locations, sites were chosen based on previous reports of guayabillo populations, either documented in the literature or through personal communications with local inhabitants. From this pre-selection we chose sites close to roads or inhabited areas, since more remote locations in the Galapagos Islands are inaccessible for sampling.

Two to five fresh leaves were taken from each sampled tree and stored in plastic bags, which were transported to the Galapagos Science Center (San Cristobal Island) for storage at −20°C. A total of 208 individuals were sampled, ranging between 4 and 34 samples per location (Table A1). We collected the greatest possible number of individuals separated by a minimum distance of 100m to minimize the possibility of sampling genetically identical individuals.

### 2.2. Molecular Methods

#### 2.2.1. Primer Design

Guayabillo specific primers for microsatellite regions were developed from a single genomic DNA extraction using the Galaxy-based bioinformatics pipeline reported by Griffiths (2013). Sequencing was performed on an Illumina MiSeq platform at the University of Manchester genomics facility using shotgun 2×250 paired-end sequencing methodology (Nextera DNA Preparation Kit, Illumina, USA). The sample used 0.33 of a flow cell and primer design was optimized for use with Platinum *Taq* DNA polymerase (Invitrogen, USA) with an optimal T_m_ of 62°C (Min 59°C, Max 62°C) and a maximum difference among primer pairs of 3°C.

A total of 2 x 1,783,686 raw sequence reads were produced, with none flagged as poor quality. Sequence length ranged from 50 to 300 bp with a reported GC content of 40%. After screening, a total of 211 primer pairs were designed to amplify SSR regions with simple motifs of 2, 3, 4 and 6 base pairs. From this list, a total of 30 loci were selected as candidates and their respective primers were synthesized; all of them target SSR loci with trinucleotide, tetranucleotide and hexanucleotide motifs (Table 1). The Tail A sequence designed and reported by Blacket et al. (2012) was added to all forward primer sequences.

**Table 1.**
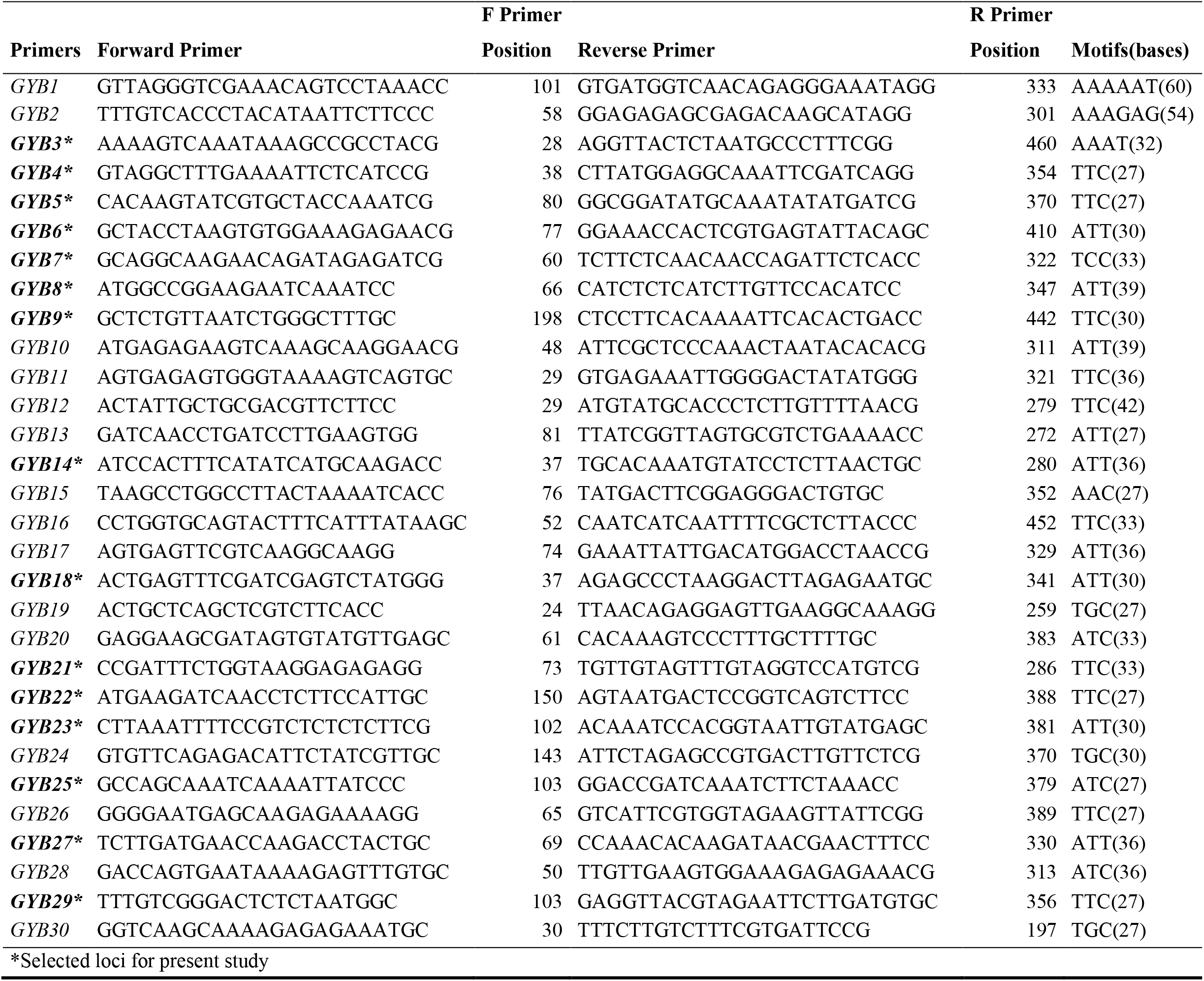
30 microsatellite loci with forward and reverse primer sequences designed for *P. galapageium.*

#### 2.2.2. DNA Extraction, Amplification and Genotyping

DNA was extracted from 25mg of leaf tissue using the CTAB method described by Saghai-Maroof et al. (1984). Isolated DNA purity and concentration were measured using a Nanodrop 1000 Spectrophotometer (Thermo Scientific, USA) and stored in TE buffer at −20°C.

After testing the 30 candidate primer sets, we selected 15 markers that amplified successfully and were polymorphic (Table 1). Amplification conditions were standardized for these 15 SSR primer sets and all samples were amplified under the following cycling conditions: 15 min at 95°C; 35 cycles of 30 sec at 94°C, 90 sec at the standardized annealing temperature, 60 sec at 72°C; 5 min at 72°C. PCR products were labeled with a fluorescent dye incorporated in the universal Tail A primer using a three-primer system (Blacket et al., 2012). Amplification products were genotyped in an ABI 313 Genetic Analyzer (Applied Biosystems) by Macrogen (Seoul, South Korea). The resulting electropherograms were analyzed using GeneMarker v. 2.4.0 (SoftGenetics LLC).

### 2.3. Statistical analyses

There is limited information regarding the ploidy of *P. galapageium*. Our observations from allele scoring suggests that up to four alleles can be found for any given locus in a single individual (see *Results*). However, 3 out of the 15 loci analyzed presented two alleles, this leads us to assume an unbalanced polyploidy which can be an indicator of allopolyploidy (Singhal et al., 1985). Furthermore, hybridization has often been observed in the *Psidium* genus (Landrum, 2017). Given these observations and the reported polyploidies in several members of the *Psidium* genus (Tuler et al., 2019), we treated *P. galapageium* as an allotetraploid and used the *polysat* package (Clark and Jasieniuk, 2011) for R (R Core Development Team, 2015) to assign alleles to different isoloci (2 isoloci per locus), thereby allowing us to process the data as diploid (Clark and Schreier, 2017). Isoloci assignment in *polysat* was performed considering all the 15 amplified SSR loci, leading to a total of 30 potential isoloci.

Null allele frequencies for each isolocus were calculated through the De Silva method (De Silva et al., 2005) implemented in *polysat.* This method requires an estimated selfing rate which is unknown in guayabillo (although the frequent occurrence of perfect flowers in this species suggests its possibility; Porter, 1968). Therefore, we used two reported selfing rates (0.5 and 0.65) from the closely related *P. guajava* (Sitther et al., 2014). Monomorphic isoloci (isolocus GYB5_2), isoloci with null allele frequencies >>0.3 given both selfing rates employed (GYB14_1, GYB14_2, GYB18_1, GYB18_2, GYB27_2), and loci that were not assigned to isoloci with an acceptable clustering quality (GYB6, GYB25) were excluded for allele frequency calculations and downstream analyses that depend on allele frequencies. Thus, from the 15 SSR loci originally amplified, we used 13 loci (from which we derived 20 informative isoloci) to describe the population genetics of our data set.

GenoDive (Meirmans and Van Tienderen, 2004) was used to determine if the analyzed guayabillo populations deviated from Hardy-Weinberg Equilibrium (HWE). The p-values obtained from the HWE test were corrected using the B-Y correction.

We used the *adegenet* package in R (Jombart and Ahmed, 2011) to determine the total number of alleles for each isoloci. Private alleles were calculated with the *poppr* package (Kamvar et al., 2014), and allelic frequencies were obtained with *polysat* using the “simpleFreq” function. Allelic richness corrected through rarefaction for different sample sizes was performed with the *basicStats* function from the diveRsity package, assuming 35 individuals sampled for all populations (Keenan et al., 2013). Significant differences among the allelic richness of different island populations were assessed with Kruskal-Wallis and Pairwise Wilcoxon tests. The same *polysat* package was used to calculate the observed and expected heterozygosity, PIC, Lynch distances and pairwise F_ST_ between islands and between sampling locations on each island, from matrixes created after isoloci reassignments. For assessing inbreeding in guayabillo, we calculated F_IS_ for each population using the calculated Ho and He values. Pairwise F_ST_ between clusters found when we conducted PCoA and STRUCTURE analyses (below) were also estimated.

We also performed an analysis of molecular variance (AMOVA) to evaluate the population differentiation between island populations in GenAlEx (Peakall and Smouse, 2012), encoding all 15 SSR markers as binary data. A Principal Coordinates Analysis (PCoA) based on Lynch distances was also plotted using *ggplot2* (Wickham, 2009).

We performed an analysis of population structure using the STRUCTURE software (Pritchard et al., 2000) following the parameters described in Meirmans et al. (2018) for dealing with polyploid data. We estimated the population structure for both the complete data set and for each island individually, using the same parameters. We evaluated between 1 and 10 potential genetic clusters (*K*) and performed 10 independent replicates for each *K* value, consisting of 1×10^6^ MCMC steps with a 1×10^5^-step burn-in period. Structure Harvester software (Earl and von Holdt, 2012) was employed to evaluate the optimum value of *K* using the Evanno method (Evanno et al., 2005). We used CLUMPP to estimate individual membership coefficients (Jakobsson and Rosenberg, 2007), and plotted them using the DISTRUCT software (Rosenberg, 2004). A plot of the relative migration levels between the three island populations was obtained by applying the Sundqvist et al. (2016) method implemented in the *divMigrate* function from the diveRsity package (Keenan et al., 2013).

Due to differences in the number of samples obtained from each island, we created subsamples for Isabela and Santa Cruz to match the San Cristobal sample size. To do so, we selected 35 individuals from Isabela and 35 from Santa Cruz (we included one random sample from each location) After this systematic downsampling, we repeated all the previously described analyses.

We used the following method in order to assigning and detecting clones: we calculated genetic distances assuming asexual reproduction under the SMM, as implemented in the GenoDive software. Missing data and unknown allele dosage were ignored. The genetic distance threshold used to classify individuals as clones (a distance of 7.0) was determined using the method suggested by Rogstad et al. (2002); note this threshold should not be equal to 0 due to the fact that mutations and genotyping errors may make identical individuals have slightly different genotypes (Meirmans and Van Tienderen, 2004). Specific clones per island were obtained. A test of clonal diversity was performed using Nei’s corrected genetic distance as summary statistic, using 999 permutations and sorting alleles over individuals within populations. Finally, clonal diversity statistics were calculated in GenoDive and bootstrap tests were performed to detect significant differences among shc (sample size-corrected Shannon index values) in different islands (999 permutations); *p*-values were corrected using the B-Y method. Clonal richness (R) was also calculated for each island and overall, as R = (G-1)/(N-1) where G is the number of genotypes detected under the established genetic distance threshold, and N is the total number of samples.

## 3. Results

### 3.1. Marker information and genetic diversity

All 208 individuals in our sample set were genotyped and included in our analyses. All of the original 15 markers amplified deviated from HWE after B-Y correction, except for locus GYB25 in the Isabela population, and GYB04 in the Santa Cruz population. The information content for the 13 markers used for data analysis (parsed as 20 isoloci) was measured through their Polymorphic Information Content indices (PIC) and ranged between 0.006 and 0.808 (only two markers showed PIC values under 0.3), with low inferred null allele frequencies (with the exception of the excluded isoloci described in Materials and Methods; Table A2). Various descriptors of genetic diversity were estimated for the populations of each of the three sampled islands (Table 2), showing similar patterns between the populations of Isabela and Santa Cruz. Compared to these, the San Cristobal Island population shows a smaller number of alleles, private alleles and both observed and expected heterozygosities (HO and HE respectively), with a higher F_ST_ fixation index. While a smaller sample size for San Cristobal (N=35, compared to N=86 in Isabela and N=87 in Santa Cruz) may account for some of these lower observed values, our downsampled analyses (i.e. reducing the samples from Isabela and Santa Cruz to maintain a constant sample size for all three; see Materials and Methods and Table A3) show a consistent reduction in the numbers of alleles (A) and private alleles (PA) for the Santa Cruz and Isabela populations, and a reduction in the observed heterozygosity for the Isabela population, but not sufficient to match the San Cristobal population H_E_ estimates; this supports our finding of a lower genetic diversity on this island. The same trend is maintained when assessing allelic richness (AR) corrected thought rarefaction among the three island populations, with a higher richness in Isabela, followed by Santa Cruz and finally San Cristobal. The difference in AR was significant between Isabela and the other two islands, both Santa Cruz (B-Y corrected Pairwise Wilcoxon test, *p* = 0.045) and San Cristobal (*p* = 0.026); nevertheless, rarefaction-corrected AR did not show significant differences among Santa Cruz and San Cristobal. Inbreeding coefficients (F_IS_) were high for the three island populations (especially Isabela and Santa Cruz) and overall for the whole dataset.

**Table 2.** Genetic diversity information of the analyzed *Psidium galapageium* populations from Isabela, Santa Cruz and San Cristobal islands: Number of individuals genotyped from each island (N), number of alleles found (A), number of private alleles (PA), rarefaction-corrected allelic richness (AR), observed heterozygosity (H_O_), expected heterozygosity/gene diversity (H_E_), F_ST_ global value for each island population, and inbreeding coefficient (F_IS_). Overall results along the three islands are also shown.

| Island | N | A* | PA* | AR<sup>s</sup> | $H_o$<sup>a</sup> | $H_E$<sup>a</sup> | $F_{ST}$ | $F_{IS}$ |
| --- | --- | --- | --- | --- | --- | --- | --- | --- |
| **Isabela** | 86 | 142 (89) | 77 (35) | 12.29 | 0.147 | 0.570 | -0.173 | 0.742 |
| **Santa Cruz** | 87 | 105 (67) | 26 (8) | 9.97 | 0.157 | 0.426 | 0.085 | 0.631 |
| **San Cristobal** | 35 | 70 (60) | 5 (1) | 8.20 | 0.119 | 0.275 | 0.286 | 0.567 |
| **Overall** | 208 | 174 | - | 10.15 | 0.141 | 0.482 | 0.230 | 0.708 |

\* Values between brackets are the number of alleles or private alleles with a frequency >0.05 within the corresponding island population.
<sup>a</sup>Indicates average across all the SSR loci analyzed.
<sup>s</sup>Standardized through rarefaction for N=35

### 3.2. Genetic differentiation of guayabillo populations

The genetic differentiation between islands, evaluated through pairwise F_ST_ genetic distances, shows a greater divergence between the San Cristobal and Santa Cruz populations, while Isabela remains equally divergent from both (Table 3). This pattern is observed with both the full and reduced data with normalized sample sizes (Table A4). Furthermore, the clustering of individuals based on Lynch genetic distances shows that the individuals from Santa Cruz are represented by two groups: a first group clearly separated from all the rest of individuals (henceforth referred to as Santa Cruz 1; Fig. 2, Fig. A1), and a second group clustering closely with individuals from Isabela and San Cristobal (henceforth referred to as Santa Cruz 2; Fig. 2, Fig. A1). This second group includes individuals from three different locations on Santa Cruz: Granillo Rojo, Garrapatero and Bellavista (Fig. A2). It should be noted, however, that the degree of population differentiation between islands appears to be limited: an AMOVA reveals that the majority of the genetic variation (72%) occurs within populations, and 28% of the variation occurs between Isabela, San Cristobal and Santa Cruz (Table 4).

**Fig. 2.**
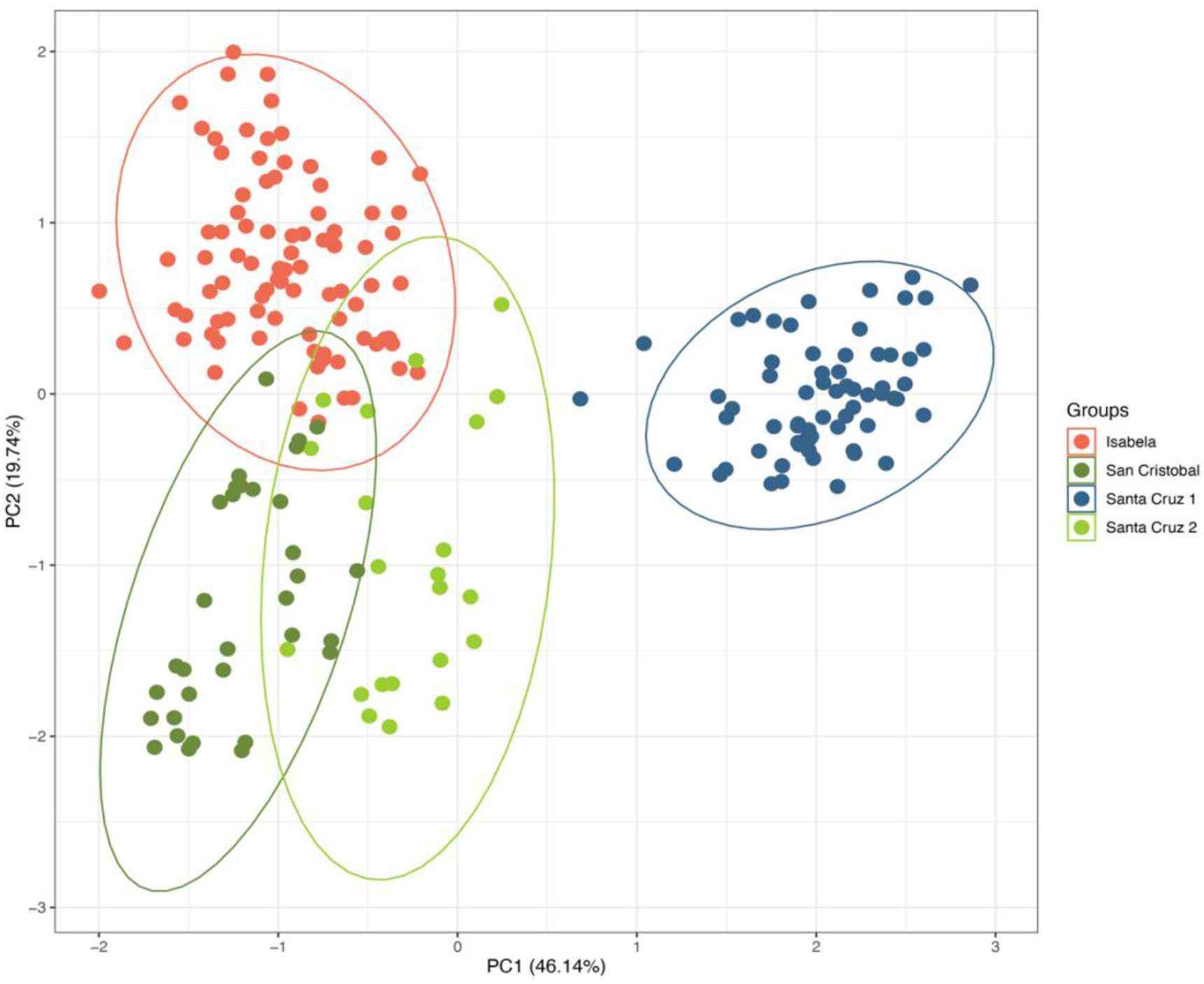
PCoA based on the Lynch distances found between the *Psidium galapageium* individuals sampled in the three islands: Isabela, San Cristobal and Santa Cruz. For Santa Cruz, both genetic clusters are indicated (Santa Cruz 1 and Santa Cruz 2).

**Table 3.**
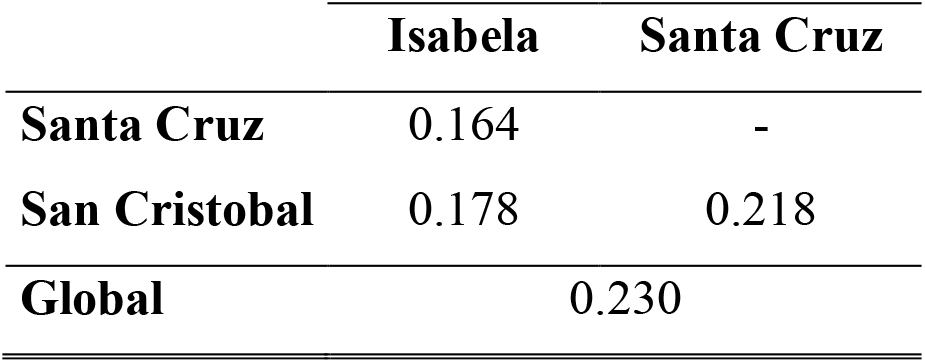
Pairwise and global F_ST_ values between the *Psidium galapageium* populations from the three islands.

**Table 4.** Analysis of molecular variance (AMOVA) between the three island populations of *Psidium galapageium*.

| Source | DF | SS | MS | Est. Var. | % |
| --- | --- | --- | --- | --- | --- |
| Among Pops | 2 | 691.79 | 345.90 | 5.12 | 28% |
| Within Pops | 205 | 2653.82 | 12.945 | 12.95 | 72% |
| Total | 207 | 3345.62 | - | 18.06 | 100% |

### 3.3. Population structure

To explore the population structure under an admixture model, the assignment coefficients for all individuals were estimated for different numbers of putative lineages, revealing higher similarities between individuals from Isabela and San Cristobal, in concordance with the clustering by genetic distances. The individuals from Santa Cruz display a greater contribution from a separate genetic stock, with some individuals showing similarities to the Isabela and San Cristobal populations (Fig. 3a). An evaluation of the optimum number of clusters that fit the data suggests that three putative lineages are observable in our data (*K*=3; Δ*K*=289.55), which highlight a closer resemblance between the genetic composition of the Santa Cruz outliers and the San Cristobal population (Fig. 3b). Overall, the three genetic lineages are determined by island, as would be expected given the physical separation and isolation between these populations. A similar analysis with the downsampled data (Fig. A3) reveals no observable differences when compared to the full data set.

**Fig. 3.**
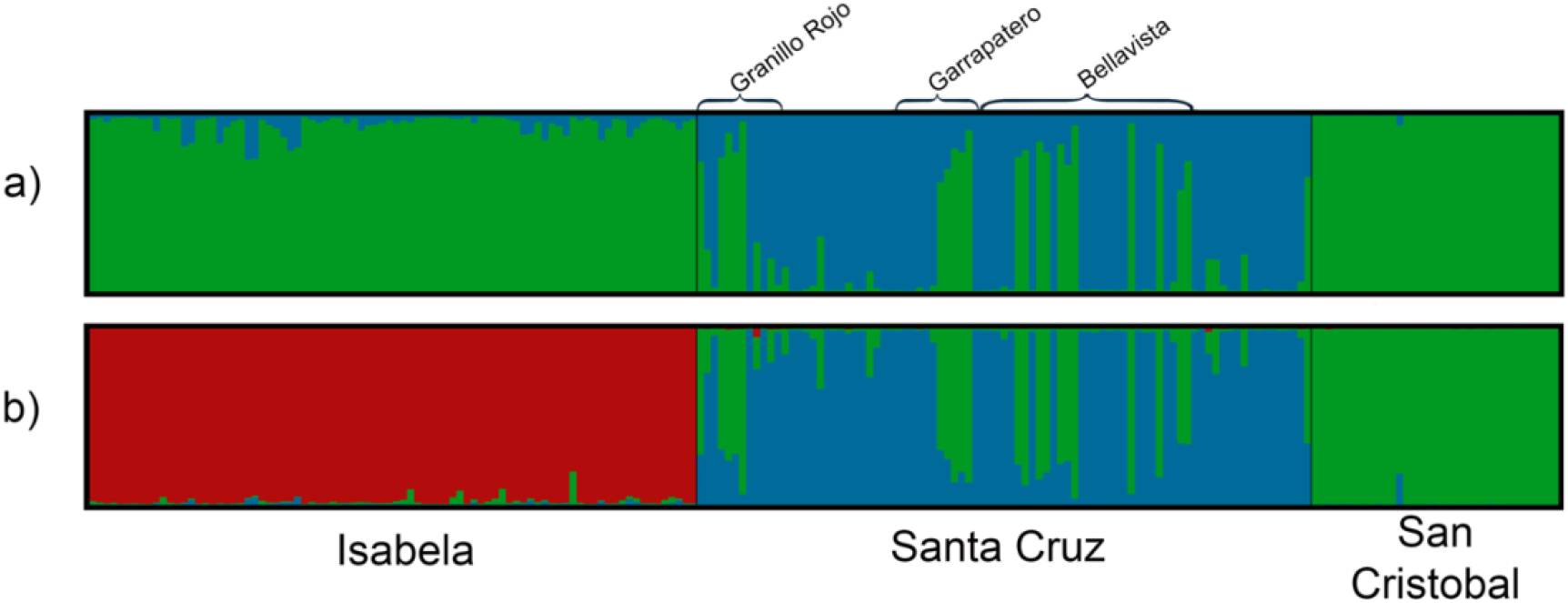
Results of the Bayesian analysis of population structure (Software STRUCTURE) under the Admixture model. The results are indicated for a) K=2 (ΔK = 134.51), and b) K = 3 which is the optimum K value (ΔK = 289.55). These values of K correspond to the clusters or lineages (represented by different colors) in which are grouped the *Psidium galapageium* individuals sampled in Isabela, Santa Cruz and San Cristobal islands. The Santa Cruz sampling sites of Granillo Rojo, Garrapatero and Bellavista (which mostly harbor individuals from the Santa Cruz 2 cluster) are marked as well.

We noted that the groupings observed through Bayesian inference in STRUCUTRE and the clusters observed in the PCoA were equivalent, reliably defining the main genetic groups in the guayabillo populations of the three islands. Pairwise F_ST_ values were calculated among these genetic groups, considering each island population individually, and including the Santa Cruz 1 and Santa Cruz 2 groups as separated entities as well. Here, the highest genetic differentiation was detected among the Santa Cruz 1 population and the populations of the other two islands: Isabela and San Cristobal. Furthermore, an important genetic differentiation was observed between the two Santa Cruz groups, comparable even to the values found among populations from different islands (Table A5).

Bayesian population structure analyses were conducted for each island. When analyzing the Isabela and Santa Cruz populations, no distinguishable population structure within each island was observed, suggesting widespread gene flow and an ancient shared history within each island (Figs. A4 and A5, respectively). The optimum K value (K=2; Fig. A5a) shows two lineages in Santa Cruz island, matching the Santa Cruz 1 and Santa Cruz 2 groups found in the PCoA (Fig. 2); however, this pattern is less clear at higher K values (Fig A5 b-d). Finally, a more distinguishable structure is observed in San Cristobal at K=2 and above, with individuals from any given sampling location tending to share their genetic background (Fig. A6).

Although limited, some migration could exist between the guayabillo populations from different islands. The relative migration analysis showed that most of the gene flow is directed towards Santa Cruz from both Isabela and San Cristobal. Outgoing migration from Santa Cruz and among Isabela and San Cristobal appears less prevalent, representing approximately half of that observed towards Santa Cruz (Fig. 4).

**Fig. 4.**
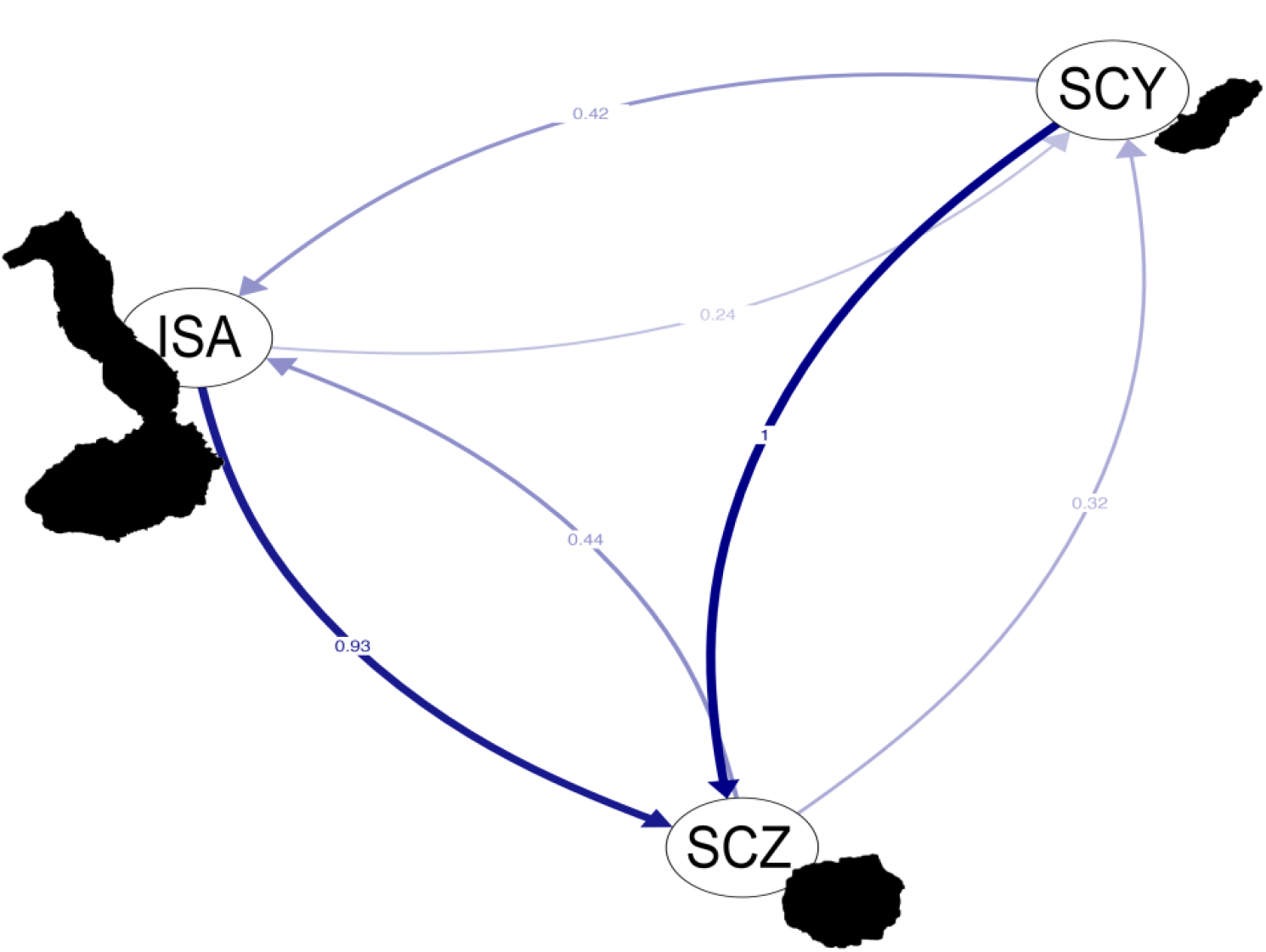
Relative migration among the guayabillo populations from Isabela (ISA), Santa Cruz (SCZ) and San Cristobal (SCY) islands.

### 3.4. Clonal assignments and clonal diversity in guayabillo

A total of 201 different unique multilocus genotypes were identified in our dataset, and 11 of the 208 analyzed guayabillo individuals (5.28%) were identified as clones of another individual; a lower number of unique genotypes was obtained when taking into account the effective number of genotypes, nonetheless they are still considerable when taking into account the total number of individuals analyzed. Clonal richness and sch values over the three populations were relatively high in general terms. Nevertheless, the San Cristobal population had the highest number of individuals sharing the same multilocus genotype (with up to five individuals having the same genotype in one case); individuals assigned to the same clone in San Cristobal belonged to different sampling locations. On the other hand, only two individuals with the same genotype were found in Isabela, as well as in Santa Cruz (coexisting in the same sampling location in both cases). Similarly, the Isabela and Santa Cruz populations showed higher clonal richness and shc values than San Cristobal (Table 5); these differences in shc were significant (Isabela vs. San Cristobal: *p* = 0.003; Santa Cruz vs. San Cristobal: *p* = 0.003).

**Table 5.** Clonal diversity statistics for the three studied island populations, and overall values: Number of individuals genotyped (N), number of clones or unique genotypes detected under the established genetic distance threshold (G), clonal richness (R) and Shannon diversity index for genotypes, corrected for sample size (shc). Calculations were performed twice: using the SSR genotyping directly without allele dosage correction for polyploids, and then using the genotypes corrected for allele dosage in polyploids.

| Island | N | G (eff)* | R | shc |
| --- | --- | --- | --- | --- |
| **Isabela** | 86 | 85 (84.0) | 0.988 | 3.564 |
| **Santa Cruz** | 87 | 86 (85.0) | 0.988 | 3.574 |
| **San Cristobal** | 35 | 30 (21.5) | 0.853 | 2.111 |
| **Overall** | 208 | 201 (190.5) | 0.966 | 3.083 |

\*Values between brackets correspond to the effective number of genotypes (G).

Finally, we found low statistical support for the hypothesis that the observed clonal diversity is explained by random mating in the three populations (Isabela: *p*=0.007; Santa Cruz: *p*=0.001; San Cristobal: *p*=0.001). This suggests that the observed clonal diversity patterns are not due to sexual reproduction; therefore, the occurrence of the same multilocus genotype in more than one individual is explained more likely by clonal or asexual reproduction rather than by sexual reproduction among related individuals.

## 4. Discussion

### 4.1. Genetic diversity in guayabillo and its contributing factors

The overall genetic diversity of 0.482 found in guayabillo is comparatively high for an insular species (Table 2). In comparison, other endemic plant species from the Galapagos such as *Scalesia affinis* (Nielsen, 2004), *Solanum galapagense* (Pailles et al., 2017), and the *Opuntia* cacti (Helsen et al., 2009) present low genetic diversity (assessed through AFLP markers, DArTSeq, and chloroplast and nuclear sequences, respectively). Similarly, other insular species outside the Galapagos *-Robinsonia evenia* and *R. gracilis* from the Juan Fernandez archipelago*-* show lower H_E_ values (0.253 and 0.345, respectively) when analyzed with SSR markers (Takayama et al., 2013). All these low diversity levels have previously been explained as a consequence of founder effects, population bottlenecks and genetic drift, typical of the colonization processes and limited ecosystems associated with most islands (Frankham, 1997; Whittaker and Fernández-Palacios, 2007). This is well illustrated by the case of the endemic Galapagos cotton *Gossypium klotzschianum*, which presents a lower genetic diversity than its closest mainland relative (and potential parental species), *G. davidsonii*, from Baja California (Wendel and Percival, 1990). Nevertheless, low genetic diversity levels are not always the case for insular species. For example, the widely distributed Galapagos endemic *Gossypium darwinii* (Wendel and Percy, 1990), and even endangered species with restricted distributions such as *Galvezia leucantha* (Guzmán et al., 2016) and *Calandrina galapagosa* (Jaramillo et al., 2011), showed relatively high genetic diversity levels in the archipelago. Similarly, insular populations of *Dacrycarpus imbricatus* from the Hainan island in China have a comparable or even a higher genetic diversity than their continental counterparts (evaluated through ISSR markers; Su et al., 2010), and some endemic plant species from Hawaii display high H_E_ values of up to 0.724 (evaluated through SSR markers; Crawford et al., 2008). Note however, that different molecular techniques have been used for assessing the diversity in the species cited; moreover, numerous factors are influencing the genetic diversity levels in each one of these cases. Therefore, care should be taken when comparing diversity levels, especially among unrelated species with different natural histories (Fernández-Mazuecos et al., 2014; Guzmán et al., 2016).

The physical characteristics of the islands where guayabillo is found could be associated to this elevated genetic diversity (Stuessy et al., 2014). Larger islands with broader altitudinal ranges host greater habitat heterogeneity (MacArthur and Wilson, 1967; Buckley, 1985; Geist et al., 2014), which in turn can favor genetic variability in a wide-distributed species as it adapts to new niches (MacArthur and Wilson, 1967; Stuessy et al., 2006; Chapman et al., 2013). In fact, morphological variation among guayabillo populations along the altitudinal gradient where the species is distributed has been observed (Valdebenito, 2018). Large island surface areas also translate into a greater capacity to host bigger populations which are more resilient to genetic drift (MacArthur and Wilson, 1967; Frankham, 1997; Vellend and Geber, 2005; Costanzi and Steiffeten, 2019). In this regard, high levels of genetic diversity in the Hawaiian silverswords (Witter and Carr, 1988) and in *G. darwinii* (Wendel and Percy, 1990) were explained in part by their large population sizes. A similar scenario might be suggested for guayabillo since the highest genetic diversity was found in Isabela island (Table 2; Table A3), which is the largest and most elevated island in the Galapagos archipelago (even if we only consider the southern part of the island where guayabillo is found) (Instituto Geofísico, n/d; Charles Darwin Foundation, 2012; Geist et al., 2014); this pattern is still observed even though our sampling covered a narrow range of the total altitudinal range of the island (109-386 m.a.s.l.). On the other hand, the San Cristobal population presented the lowest genetic diversity among our sampling sites (Table 2; Table A3), coinciding with the island’s smaller size and narrower altitudinal range (Latorre, 1991; Charles Darwin Foundation, 2012); our guayabillo samples cover approximately half of this range (71-310 m.a.s.l.). Finally, Santa Cruz, where we obtained an intermediate H_E_ but not a higher AR than San Cristobal, constitutes an intermediate altitudinal and land mass range between Isabela and San Cristobal (Grenier, 2007; Charles Darwin Foundation, 2012). These general trends are not surprising and have also been observed in the Galapagos endemics *S. affinis* (Nielsen, 2004) and *G. darwinii* (Wendel and Percy, 1990), which showed greater genetic variation in Isabela compared to smaller and lower islands as Santa Cruz, Floreana (both species) and San Cristobal (*G. darwinii* only). A greater abundance and genetic diversity have also been reported for the endemic tomatoes *S. cheesmaniae* and *S. galapagense* in western islands like Isabela, something that was also attributed tentatively to the unusually high precipitation for this part of the archipelago (Rick and Fobes, 1975; Pailles et al., 2017). Knowing that plant richness is positively linked with precipitation in the tropics (Gentry, 1982), this could also explain the greater diversity observed in the Isabela guayabillo population, as well as in the case of other endemic plants. Thus, despite being one of the youngest islands in the Galapagos (Geist et al., 2014), Isabela would present certain conditions that favor diversity in endemic plant species, though other factors should also be considered when interpreting these genetic diversity patterns.

The limited available evidence suggests a complex evolutionary history for the guayabillo which may also partially explain the genetic diversity patterns observed in the species. Firstly, guayabillo could be a polyploid species, as suggested by our genotyping, where up to four different alleles were observed for several loci. Furthermore, morphological studies point to guayabillo being very phenotypically similar to the hybrids between two mainland close relatives: *Psidium oligospermum* and *P. schenckianum* (Landrum, 2017); Landrum (2017) also hypothesized that the hybrids of these two species were able to spread all over tropical America following the hybridization event, opening the possibility that they may have reached the Galapagos. With these antecedents, the Galapagos guayabillo could tentatively be an allopolyploid, with *P. oligospermum* and *P. schenckianum* as putative parental species, a hypothesis that could be confirmed through phylogenetic analyses of the *P. oligospermum* complex. In any case, hybridization is quite frequent in the *Psidium* genus (Machado-Marques et al., 2016; Landrum, 2017), and this potential allopolyploidy in guayabillo could also be one of the reasons behind its moderately high genetic diversity despite it being an insular endemic species. Allopolyploids show a tendency towards higher heterozygosity and genetic variability levels compared to diploids as they draw from the gene pools of two separate species, which might be the case if guayabillo is in fact an allopolyploid (Soltis and Soltis, 2000; Chen and Ni, 2006). Moreover, previous hybridization and allopolyploidy have been tightly associated with the success of the colonization of oceanic islands by plants (Barrier et al., 1999; Wendel and Cronn, 2003; Madlung, 2013). These ideas could also be supported by the high genetic diversity found in the widespread tetraploid *G. darwinii,* for example. However, polyploidy is not a requisite or a guarantee for high genetic diversity levels. For instance, the *Opuntia* cacti are hexaploid and still display low genetic diversity levels (Helsen et al., 2009). Likewise, there are diploid insular plant species that show moderate to high levels of genetic diversity (e.g. Crawford et al., 2008; Takayama et al., 2013; Takayama et al., 2015). Therefore, polyploidy is not the only aspect of evolutionary history that should be addressed for interpreting the genetic diversity observed in guayabillo.

The mechanism of speciation may also be important to explain the genetic diversity in insular plant populations (Stuessy et al., 2014; Takayama et al., 2015). Cladogenesis, for instance, generates several daughter species, each one with low levels of genetic diversity as observed in the endemic *Opuntia* cacti from different islands in the Galapagos archipelago (Helsen et al., 2009). On the other hand, guayabillo has not been reported to split into separate species in different islands (Porter, 1968; McMullen, 1999), this is also supported by our genetic data. Even though we observed a genetic structure between different islands and a limited inter-island gene flow (Fig. 3), we do not have evidence to claim the populations from Isabela, Santa Cruz and San Cristobal are distinct species. Firstly, the pairwise F_ST_ values among island populations, though high, are not high enough to reach that conclusion (Table 3; Table A5). Secondly, we would have expected a higher percentage of the total diversity explained by diversity among islands if they were different species (Table 4). Finally, most of the individuals, including samples from distinct islands, are clustered together in the PCoA (with the only exception of the Santa Cruz 1 group; Fig. 2). In consequence, the history of guayabillo aligns with the speciation mechanism of anagenesis, where different processes such as mutation accumulation, recombination, and local adaptation would have created more and new genetic diversity which was kept in a single species (Stuessy et al., 2006; Takayama et al., 2015). Other species in the Galapagos also continue to be a single species despite having populations separated in different islands; *G. leucantha* for example, keeps a moderate-low genetic differentiation even among islands, leading to a high genetic and morphological diversity within this species as a whole (Guzmán et al., 2016).

The reproductive biology of the species is also important to understand the genetic diversity patterns (Stuessy et al., 2014), which is poorly understood for guayabillo (Valdebenito, 2018) Complementary research on outcrossing-selfing rates, pollinization, seed dispersal and germination rates is required to determinate the effect of these factors on the genetic diversity of guayabillo. However, our genetic data could shed some light on these topics. Firstly, we found low H_O_ values compared to H_E_, as well as high F_IS_ values in all the three islands (Table 2), which suggests that inbreeding and/or selfing (guayabillo has bisexual flowers and selfing is concurrent in the *Psidium* genus) in all guayabillo populations could be prevalent (Wright, 1951; Loeschcke et al., 1994; Frankham, 1998; Sitther et al., 2014). Guayabillo is also known to reproduce through root suckers which may lead to clones (Aldaz, 2008); however, we made sure to sample physically distant individuals to avoid the collection of this type of clones. No direct studies have been performed to test other kinds of clonal reproduction in guayabillo such as apomixis. In any case, our results do show a couple of individuals from distant locations sharing the same multilocus genotype. These cases appear to be sporadic, and most of the sampled individuals represent unique genotypes (Table 5). Similarly, guayabillo presents a higher clonal diversity levels than other plants which are actually known to reproduce clonally such as *Ziziphus celata* (Gitzendanner et al., 2012) and *Trillium recurvatum* (Mandel et al., 2019). Indirect evidence of outcrossing in guayabillo was obtained through the observation of flowers being visited by the Galapagos carpenter bee *Xylocopa darwinii* (Valdebenito, 2018), also an important pollinator for several other endemic plants of the Galapagos (see Jaramillo et al., 2014; Guzmán et al., 2016). Likewise, the higher within-island diversity compared to the between-island diversity (Table 4) aligns with what would be expected for a cross-pollinating plant species, despite the non-negligible 28% of among-island diversity. In light of previous descriptions of the ecology of the species (Aldaz, 2008; Valdebenito, 2018), our genetic data suggests that guayabillo might combine different reproductive mechanisms including clonal reproduction, selfing and outcrossing. The common guava, a close relative of the guayabillo, shows similar reproductive mechanisms (Urquía et al., 2019). This could also explain the relatively high genetic diversity we found in guayabillo, as well as its success in colonizing the Galapagos archipelago. Clonal reproduction and selfing would have aided in the fast spread of the species over the islands during the first stages of the colonization, while the increasing population sizes solidified the endurance of high-fitness genotypes (Pluess and Stöcklin, 2004; Silvertown, 2008). In fact, self-compatibility would be the general rule for insular plants as it is essential for the establishment in new islands (Baker, 1955; Chamorro et al. 2012). However, this kind of reproduction is known to reduce genetic diversity and drive inbreeding depression (and its associated consequences such as disease susceptibility and low mate availability; Kwak and Bekker, 2006, Honnay and Jacquemyn, 2008), as seen in the highly autogamous endemic tomatoes *S. cheesmaniae* and *S. galapagense* (Rick, 1983; Pailles et al., 2017). Hence, thanks to its potential capacity for combining asexual reproduction and outcrossing, guayabillo would have also been able to maintain its genetic diversity and a wide variety of clones and genotypes, retaining the species’ adaptive potential while keeping the advantages of clonal spread (Ward et al., 2008). This hybrid system would be beneficial in fluctuating and unpredictable environments, characteristics that recall the nature of the Galapagos Islands (Bengtsson, 2000; Silvertown, 2008; Capotondi et al., 2015). Thus, it is not surprising that other plants in the Galapagos such as *C. galapagosa* also hold considerable levels of genetic diversity, perhaps through outcrossing (Jaramillo et al., 2011), while *Lycium minumum* developed sexual dimorphism to equilibrate self-compatibility and outcrossing (Levin et al., 2015).

A case could also be made regarding the effects of human disturbance in the genetic diversity of the species, particularly for the San Cristobal population. The three islands host permanent human populations that in some cases use guayabillo as a source of wood. Moreover, these islands also sustain agricultural areas in their highlands, where local-scale cultivation and animal husbandry activities are developed. San Cristobal contains the largest agricultural area relative to its size, occupying the majority of its humid highlands (Rivas-Torres et al., 2018), and one of the oldest permanent settlements in the archipelago established during the second half of the XIX century (Latorre, 1997; Lundh, 2004). These events could have affected the guayabillo populations disproportionately when compared to the other islands. Santa Cruz was colonized more recently by humans, producing a milder historical disturbance (Kricher, 2006; Parque Nacional Galapagos, 2016), although it currently hosts a larger human population (Instituto Nacional de Estadísticas y Censos, 2016). On the other hand, Isabela sustains the smallest agricultural area in proportion to island size (Rivas-Torres et al., 2018) and the smallest human population (Instituto Nacional de Estadísticas y Censos, 2016). A direct effect of these anthropogenic activities is the fragmentation of habitats which can lead to genetic drift (Frankham et al., 2010), endogamy and inbreeding depression (Wright, 1951; Frankham, 1998; Nielsen, 2004), and could partially explain the higher within-island F_ST_ value observed for the San Cristobal population (Table 2). Fragmented and decimated populations also experience a fast fixation of alleles, and populations within fragments risk differentiating to the point that of sexual incompatibility (Gitzendanner et al., 2012). It is also noteworthy that the San Cristobal population presented a lower clonal diversity and a higher proportion of individuals sharing the same genotype compared to the other two islands (Table 5). This could be a consequence of the depauperated genetic diversity in this population, leading to less alleles and therefore, less possible genotypes (Jaramillo et al., 2011). *S. affinis* might represent a similar scenario to guayabillo in San Cristobal: habitat loss and intensive grazing by donkeys and goats has caused widespread habitat loss in the small Floreana and Santa Cruz *S. affinis* populations leading to a reduction of the genetic diversity (Nielsen, 2004). Likewise, habitat loss and aggressive herbivory from introduced animals has fragmented and decimated the populations of *C. galapagosa* (Jaramillo et al., 2011) and *G. leucantha* (Guzmán et al., 2016).

Nonetheless, our findings present a positive outlook for guayabillo. The relatively high levels of genetic diversity found in this species suggest that these populations show some potential resilience to environmental perturbations (Reusch et al., 2005; Jump et al., 2009). An increased adaptive potential would certainly be an asset for the species in face of threats such climate change and habitat alteration associated to human activities (Adsersen et al., 1988; Whittaker, 1998; Tye et al., 2007; Dal Forno et al,, 2017); however, the survivability of any species is not determined exclusively by its genetic adaptive potential, and other factors must be better studied to understand the conservation status of guayabillo in the archipelago. For instance, the interactions between guayabillo and multiple invasive plant species in the Galapagos, particularly those found in the highlands and transitional forests such as blackberries and Cuban cedars (*Cedrela odorata*) (Sakai et al., 2001; Tye et al., 2007), remain unknown. A particular emphasis should be placed on the invasive common guava due to its close relatedness to guayabillo and the fact that they share similar distributions, life history traits, pollinators and dispersers (Blake et al., 2012; Valdebenito, 2018). The high frequency of hybridization events in the *Psidium* genus (Landrum, 2017) should also be considered, as this combination of factors might facilitate the generation of (currently unreported) interspecific hybrids (Torres and Gutiérrez, 2018). This phenomenon can lead to genetic erosion, outbreeding depression, and genetic swamping in the guayabillo (López-Caamal et al., 2014; Ellstrand and Rieseberg, 2016; Chafin et al., 2019) while enriching the currently low genetic diversity of the guava populations of the Galapagos, further enhancing its invasive potential (Urquía et al., 2019). Such a case has already been reported in an insular *Psidium* endemic, *P. socorrense,* where hybridization with an introduced close relative took place in a particular zone of Socorro Island (López-Caamal et al., 2014).

### 4.2. Population structure and connectivity between islands

The observed patterns of genetic diversity do not necessarily match the population structures in different islands. Santa Cruz is the only island where two clearly separated genetic clusters were found (Figs. 2, 3, A1 and A3), while the populations in Isabela and San Cristobal behave as a single panmictic population. One of these Santa Cruz clusters, Santa Cruz 2, was exclusively made up of individuals from sampling sites within the transition zone (Granillo Rojo), the dry lowlands (Garrapatero) and some individuals from the Bellavista site (which is closely located to Garrapatero). On the other hand, Santa Cruz 1 predominantly included individuals from the humid highlands and the agricultural zone (Figs. 3, A2 and A5). These clusters may correspond to two different guayabillo ecotypes, a more generalist ecotype (Santa Cruz 1) and a dry climate ecotype (Santa Cruz 2) adapted to the transition zones and the lowlands. Interestingly, Valdebenito (2018) observed morphological differences among guayabillo individuals from the highlands and the lowlands in San Cristobal, (monopodial trees in the highlands, smaller shrubs in the lowlands); more significantly, lowland individuals would flower earlier, which could represent a temporal reproductive barrier between them and highland individuals. Although we did not identify different genetic groups in San Cristobal as we would have expected from previous observations, it highlights the possibility of two ecotypes in Santa Cruz; phenological and morphological studies of guayabillo in this island are currently being carried out (Valdebenito, pers. comm.), and they would certainly elucidate our hypothesis. This would entail a degree of genetic differentiation (observed as a high proportion of within-population variability in the AMOVA; Table 4) and adaptation to different climatic and ecological niches, phenomena which cannot be further explored with our current data. More in-depth research into the population genetics and ecology of the species in this island is essential to determine whether the concept of an ecotype might apply to this scenario. The emergence of different ecotypes and even parapatric speciation along environmental gradients have been previously reported in plants, such as the two sister species of the genus *Senecio* distributed along different altitudes at Mount Etna in Italy which may have arisen through these mechanisms (Chapman et al., 2013; Chapman et al., 2016).

It is also interesting to point out the genetic similarities observed in the PCoA between Santa Cruz 2 and the individuals from San Cristobal and Isabela (Figs. 2, A1 and A2). This could be interpreted as a link between Santa Cruz 2 and the populations on the other islands, particularly in San Cristobal (see Fig 3b). Under this scenario, the Santa Cruz 1 lineage would have naturally diverged from the other populations on different islands (Table A5), while the Santa Cruz 2 represents a more recent introduction. Given this possibility, Santa Cruz 2 (or its ancestors) could have adapted to the drier habitats before reaching Santa Cruz, helping it to settle into its current distribution (it would be expected that, upon arriving to a new island, plants would first encounter the more arid habitats in the lowlands near the coast; Kricher, 2006; Rivas-Torres et al., 2018). If this genetic connectivity between Santa Cruz 2 and the other islands is not spurious, the previously described population structure would be better explained by this rationale rather than a local adaptation to different environments, or through a combination of both scenarios. Note that Santa Cruz 1 appears surprisingly distinct, even compared to other individuals of Santa Cruz (Figs. 2 and A1; Table A5). Before, we supported the unification of guayabillo as a single species (Section 4.1), and this seems to be true even for this separated group, since it still maintains some (limited) gene flow with the rest of the Santa Cruz populations as seen in the STRUCTURE analysis (Fig. 3) and the pairwise F_ST_ values (Table A5). Nevertheless, the differentiation among the Santa Cruz 1 and Santa Cruz 2 groups is equivalent to the differentiation seen among different islands, and likewise, Santa Cruz 1 is the genetic group with the highest inter-island differentiation seen in guayabillo (Table A5). Therefore, this leads to either a strong (potentially early) divergence of the Santa Cruz 1 group from the rest of the species, or the possibility of two different colonization events of the ancestral guayabillo into the archipelago an alternative hypothesis. The latter has been as proposed for another Galapagos endemic, *Croton scouleri,* which also displays a notable genetic and morphological variability (Rumeu et al., 2016). The current data is limited and these hypotheses remain speculative, certainly a broader sampling range across the archipelago and the use of more powerful molecular markers are necessary to solve the ancestry relations among different populations and lineages from different islands.

The degree of gene flow between islands is a key factor in explaining the previously described population structure. On a broader scale, there’s a clear genetic differentiation between the populations of the three islands, made evident by the high pairwise F_ST_ values observed (Table 3, Table A4) and by individuals clearly clustering according to their island of origin (Figs. 2, 3b, A1 and A3b). Furthermore, a good proportion of the alleles found in each guayabillo population were private alleles (Table 2), highlighting the independent evolutionary histories on each island. Selfing, inbreeding and clonal reproduction (to a lesser extent) in each island population would have led to the fast fixation of distinct alleles that, together with new mutations, could contribute to the current genetic structure and population differentiation (Rick, 1983; Hamrick and Godt, 1996). Moreover, this degree of differentiation suggests a limited gene flow between islands, similar to other endemic species such as *S. affinis* and the *Opuntia* cacti (Nielsen, 2004; Helsen et al., 2009). The oceanic waters that separate the islands are evidently an important barrier for inter-island gene flow in guayabillo and other endemic plants of the Galapagos, especially considering that its fruits and seeds are unlikely to be dispersed through long distances over the ocean (Porter, 1968; Porter, 1976; Ward and Brookfield, 1992; McMullen, 1999). In addition, none of the known animal dispersers of guayabillo seeds -Giant Tortoises, and possibly small passerine birds (Blake et al., 2012; Guerrero and Tye, 2009; Heleno et al., 2013)-would frequently cross large expanses of ocean among islands (Petren et al., 2005; Gerlach et al., 2006; Smith, 2009). Nevertheless, we cannot exclude the possibility of occasional gene flow between guayabillo populations on different islands, potentially mediated by human beings transporting seeds or propagative material between islands as a trading activity (Wiggins et al., 1971), or by widespread and generalist pollinators like *X. darwini* which are also strong flyers that can be easily carried over the ocean by the wind (McMullen, 1990; Smith, 2009; Traveset et al., 2013; Valdebenito, 2018). In fact, our migration analysis shows that most of the limited inter-island migration is directed towards Santa Cruz, in the center of the archipelago (Fig. 4), matching the confluence of sea currents and winds acting upon the Galapagos (Merlen, 2014). Note also that this gene flow to Santa Cruz may also explain the presence of the Santa Cruz 2 group and its close relationship with the populations of the other two islands (Figs. 2 and 3). Despite the notoriety of the oceanic barrier, other plants as *L. minimum* (where a significant population structure among islands was also found; Levin et al., 2015) or *G. leucantha* (Guzmán et al., 2016) are also able to hold some inter-island gene flow, which has been attributed respectively to the action of bird dispersers and the long-range pollination by *X. darwinii*.

In other endemic plant species of the Galapagos, including *S. cheesmaniae* (Pailles et al., 2017), *L. minimum* (Levin et al., 2015) and *G. darwinii* (Wendel and Percy, 1990), a clear genetic structure pattern separating populations of the western and eastern islands was observed. Such pattern apparently follows the progression rule, separating populations from older and younger islands and suggesting an east-west migration (from old to young islands) following the movement of the archipelago with the Nazca plate (Geist et al., 2014; Merlen, 2014; Levin et al., 2015; Pailles et al., 2017). However, the natural history of guayabillo appears more complicated than that. Putting aside the possibility of a second introduction of guayabillo into Santa Cruz, we would expect a greater genetic similarity between the closer islands (both temporally and geographically), a pattern that doesn’t hold true given the closer relation between the populations from Isabela and San Cristobal compared to the Santa Cruz individuals (Figs. 2 and 3a; Geist et al., 2014). The lack of an evident clustering of individuals from older and younger islands appears to refute the progression rule for guayabillo in the sampled islands. Note however that the compact spatial clustering of the archipelago in two-dimensional space (Geist et al., 2014; Shaw and Gillespie, 2016) make this observation not surprising. The ancestors of guayabillo, as several other endemic species, have not always moved progressively from older to younger islands, instead moving through one or more of thousands of alternative pathways for spreading over the archipelago beginning from a single island (Geist et al., 2014). Hence, a movement of guayabillo from Isabela to San Cristobal or vice-versa, is perfectly possible. The majority of the Galapagos endemic species, especially the most vagile organisms, did not follow the progression rule during their colonization (Shaw and Gillespie, 2016), including the endemic *Opuntia* cacti (where individuals from Isabela were contained in the same clade as the individuals of the oldest islands, Española and San Cristobal; Helsen et al., 2009) and several animals such as giant tortoises (Caccone et al., 2002), Darwin finches (Grant and Grant, 2008), land iguanas (Gentile et al., 2009), and various insect taxa (Schmitz et al., 2007; Sequeira et al., 2008). There are many other possibilities behind the biogeographic history of guayabillo, a task that could be better addressed through phylogenetic analyses using appropriate markers, and with the inclusion of samples from all the islands where guayabillo is distributed.

## 5. Conclusions

Our current data highlights some of the key questions that can be postulated about the history, evolution and future prospects of the guayabillo on the Galapagos Islands. Its relatively high genetic diversity, unusual for an insular species, could suggest an ancient history and extensive opportunities to differentiate through isolation from neighboring islands or through adaptation to new microclimates and niches. Several aspects would be promoting this genetic variability in guayabillo, including its potential allopolyploid origin followed by anagenesis, and its capacity of holding outcrossing together with selfing and clonal reproduction; bigger and higher islands with less human impact as Isabela, also would be capable of harboring more genetic diversity on them. The relatively well-defined population structure we found in guayabillo between different islands, may also be reflecting the effects of reproductive mechanisms and oceanic barriers on the spread of this species, shedding some light into the main drivers of its range and mobility. However, finer details like a weak yet discernible differentiation process within Santa Cruz raise multiple hypotheses about the adaptive processes or potential gene flow between islands. It is likely that a combination of factors drives the population dynamics of guayabillo on the Galapagos, and the relatively recent human presence may play a more important role in its future.

Our results provide, for the first time, an insight into the population genetics of the species while emphasizing the importance of using genetic tools to better understand the natural history of a species. Likewise, this genetic data can be informative for the implementation of conservation strategies for the species. For instance, our data suggests that the San Cristobal population could be the most vulnerable among the ones analyzed in this study, prioritizing the implementation of management actions in this island. The possible fragmentation issue and its lower clonal diversity could be one of the biggest concerns in this case, since this may lead to more diversity loss due to genetic drift, and mate incompatibility among subpopulations (Scobie and Wilcock, 2009; Gitzendanner et al., 2012). Thus, multi-genotype populations should be promoted and established in this island, for example by translocating or outcrossing individuals from different fragments or by allowing corridors in the farming zone of San Cristobal to favor gene flow (Gitzendanner et al., 2012). The Isabela population on the other hand, thanks in part due to the lower human impact and big dimensions of the island, appears to harbor the highest genetic variability in the studied islands, making it a potential germplasm reservoir for the species. It must be also considered also that the populations of each island represent unique gene pools, and in particular Santa Cruz, counts on two very different genetic lineages (potentially different ecotypes). These genetic clusters need to be considered independently for conservation purposes and for ex-situ collections and potential breeding programs (Gitzendanner et al., 2012; Jaramillo et al., 2011; Guzmán et al., 2016). Note that maximizing genetic diversity is essential for restoring endangered plant species, as has already been observed with in the successful recovery of *C. galapagosa* in San Cristobal Island (Jaramillo et al., 2011). Finally, a holistic conservation approach is necessary in the Galapagos as well, not only to protect guayabillo but all its flora and fauna as well (Atkinson et al., 2008; Carrion et al., 2011; DPNG, 2016). Finally, as basic biology questions (such as the ploidy of the species) are answered and new tools (such as genomic analysis pipelines) are developed, the current understanding of this endemic plant may be further developed and applied to its adequate conservation.

## Acknowledgements

We would like to thank Ricardo Campoverde, Carolina Cazco, Andrea Soria, Liseth Salazar, Gabriela Bruque and Sara Ponce for their contributions to the experimental phase of this project. We are grateful to Hugo Valdebenito, María José Pozo, Marcelo Loyola, Juan Delgado, Viviana Jaramillo and Daniel García for their assistance during the field work. We would also like to thank Hugo Valdebenito (USFQ), Todd Vision and Bryan Reatini (UNC) for the valuable conversations about the guayabillo throughout the execution of this project. We are also grateful for the support provided during the course of this investigation by the Galapagos Science Center staff and the Galapagos National Park.

In accordance with Ecuadorian regulations, plant material was obtained under the Genetic Resources Permit No. MAE-DNB-CM-2016-0041, granted by the Ministerio del Ambiente Ecuador to Universidad San Francisco de Quito.

## Funding

This study was supported by the Galapagos Science Center and Universidad San Francisco de Quito.

**Appendix A. Supplementary data**

## References

Adsersen, A., Adsersen, H., Brimer, L., 1988. Cyanogenic constituents in plants from the Galápagos Islands. Biochemical Systematics and Ecology 16, 65–77. https://doi.org/10.1016/0305-1978(88)90120-2

Aldaz, I., 2008. Manual de especies nativas y endémicas de Galápagos. Editorial FLACSO. https://biblio.flacsoandes.edu.ec/libros/digital/54629.pdf (In Spanish)

Atkinson, R., Rentería, J. L., Simbaña, W., 2008. The consequences of herbivore eradication on Santiago: are we in time to prevent ecosystem degradation gain? (Galapagos Report 2007–2008). CDF, GNP and INGALA, Puerto Ayora, Galápagos, Ecuador, 121–124.

Baker, H.G., 1955. Self-compatibility and establishment after “longdistance” dispersal. Evolution 9, 347–349.

Barrier, M., Baldwin, B.G., Robichaux, R.H., Purugganan, M.D., 1999. Interspecific hybrid ancestry of a plant adaptive radiation: allopolyploidy of the Hawaiian silversword alliance (Asteraceae) inferred from floral homeotic gene duplications. Mol. Biol. Evol. 16, 1105–1113. https://doi.org/10.1093/oxfordjournals.molbev.a026200

Bengtsson, C., 2000. The balance between sexual and asexual reproduction in plants living in variable environments. Journal of Evolutionary Biology 13, 415–422. https://doi.org/10.1046/j.1420-9101.2000.00187.x

Bensted-Smith, R., 2002. A biodiversity vision for the Galapagos Islands. Charles Darwin foundation and world wildlife fund.

Blacket, M.J., Robin, C., Good, R.T., Lee, S.F., Miller, A.D., 2012. Universal primers for fluorescent labelling of PCR fragments--an efficient and cost-effective approach to genotyping by fluorescence. Mol Ecol Resour 12, 456–463. https://doi.org/10.1111/j.1755-0998.2011.03104.x

Blake, S., Wikelski, M., Cabrera, F., Guezou, A., Silva, M., Sadeghayobi, E., Yackulic, C.B., Jaramillo, P., 2012. Seed dispersal by Galápagos tortoises. Journal of Biogeography 39, 1961–1972. https://doi.org/10.1111/j.1365-2699.2011.02672.x

Buckley, R.C., 1985. Distinguishing the Effects of Area and Habitat Type on Island Plant Species Richness by Separating Floristic Elements and Substrate Types and Controlling for Island Isolation. Journal of Biogeography 12, 527–535. https://doi.org/10.2307/2844908

Caccone, A., Gentile, G., Gibbs, J.P., Frirts, T.H., Snell, H.L., Betts, J., Powell, J.R., 2002. Phylogeography and history of giant Galápagos tortoises. Evolution 56, 2052–2066. https://doi.org/10.1111/j.0014-3820.2002.tb00131.x

Capotondi, A., Wittenberg, A.T., Newman, M., Di Lorenzo, E., Yu, J.-Y., Braconnot, P., Cole, J., Dewitte, B., Giese, B., Guilyardi, E., Jin, F.-F., Karnauskas, K., Kirtman, B., Lee, T., Schneider, N., Xue, Y., Yeh, S.-W., 2015. Understanding ENSO Diversity. Bull. Amer. Meteor. Soc. 96, 921–938. https://doi.org/10.1175/BAMS-D-13-00117.1

Carlquist, S., 1974. Island biology. Columbia University Press, New York.

Carrion, V., Donlan, C.J., Campbell, K.J., Lavoie, C., Cruz, F., 2011. Archipelago-Wide Island Restoration in the Galápagos Islands: Reducing Costs of Invasive Mammal Eradication Programs and Reinvasion Risk. PLoS One 6. https://doi.org/10.1371/journal.pone.0018835

Chafin, T.K., Douglas, M.R., Martin, B.T., Douglas, M.E., 2019. Hybridization drives genetic erosion in sympatric desert fishes of western North America. Heredity 123, 759–773. https://doi.org/10.1038/s41437-019-0259-2

Chamorro, S., Heleno, R., Olesen, J.M., McMullen, C.K., Traveset, A., 2012. Pollination patterns and plant breeding systems in the Galapagos: a review. Ann. Bot. 110, 1489–1501. https://doi.org/10.1093/aob/mcs132

Chapman, M.A., Hiscock, S.J., Filatov, D.A., 2013. Genomic divergence during speciation driven by adaptation to altitude. Mol. Biol. Evol. 30, 2553–2567. https://doi.org/10.1093/molbev/mst168

Chapman, M.A., Hiscock, S.J., Filatov, D.A., 2016. The genomic bases of morphological divergence and reproductive isolation driven by ecological speciation in Senecio (Asteraceae). J. Evol. Biol. 29, 98–113. https://doi.org/10.1111/jeb.12765

Charles Darwin Foundation, 2012. A biodiversity vision for the Galapagos Islands. In: Bensted-Smith, R. (Ed.), A biodiversity vision for the Galapagos Islands. CDF.

Chen, J.Z., Ni, Z., 2006. Mechanisms of genomic rearrangements and gene expression changes in plant polyploids. Bioessays 28, 240–252. https://doi.org/10.1002/bies.20374

Christie, D.M., Duncan, R.A., McBirney, A.R., Richards, M.A., White, W.M., Harpp, K.S., Fox, C.G., 1992. Drowned islands downstream from the Galapagos hotspot imply extended speciation times. Nature 355, 246–248. https://doi.org/10.1038/355246a0

Cimadom, A., Causton, C., Cha, D.H., Damiens, D., Fessl, B., Hood-Nowotny, R., Lincango, P., Mieles, A.E., Nemeth, E., Semler, E.M., Teale, S.A., Tebbich, S., 2016. Darwin’s finches treat their feathers with a natural repellent. Sci Rep 6, 34559. https://doi.org/10.1038/srep34559

Clark, L.V., Jasieniuk, M., 2011. POLYSAT: an R package for polyploid microsatellite analysis. Mol Ecol Resour 11, 562–566. https://doi.org/10.1111/j.1755-0998.2011.02985.x

Clark, L.V., Schreier, A.D., 2017. Resolving microsatellite genotype ambiguity in populations of allopolyploid and diploidized autopolyploid organisms using negative correlations between allelic variables. Mol Ecol Resour 17, 1090–1103. https://doi.org/10.1111/1755-0998.12639

Costanzi, J.-M., Steifetten, Ø., 2019. Island biogeography theory explains the genetic diversity of a fragmented rock ptarmigan *(Lagopus muta)* population. Ecology and Evolution 9, 3837–3849. https://doi.org/10.1002/ece3.5007

Crawford, K.M., Whitney, K.D., 2010. Population genetic diversity influences colonization success. Mol. Ecol. 19, 1253–1263. https://doi.org/10.1111/j.1365-294X.2010.04550.x

Crawford, N.G., Hagen, C., Sahli, H.F., Stacy, E.A., Glenn, T.C., 2008. PERMANENT GENETIC RESOURCES: Fifteen polymorphic microsatellite DNA loci from Hawaii’s *Metrosideros polymorpha* (Myrtaceae: Myrtales), a model species for ecology and evolution. Mol Ecol Resour 8, 308–310. https://doi.org/10.1111/j.1471-8286.2007.01937.x

Dal Forno, M., Bungartz, F., Yanez-Ayabaca, A., Lacking, R., Lawrey, J. D., 2017. High levels of endemism among Galapagos basidiolichens. Fungal diversity 85, 45. https://doi.org/10.1007/s13225-017-0380-6

De Silva, H.N., Hall, A.J., Rikkerink, E., McNeilage, M.A., Fraser, L.G., 2005. Estimation of allele frequencies in polyploids under certain patterns of inheritance. Heredity (Edinb) 95, 327–334. https://doi.org/10.1038/sj.hdy.6800728

Dirección del Parque Nacional Galápagos (DPNG), 2016. Proyecto de control y erradicación de especies invasoras prioritarias para la reducción de la vulnerabilidad de especies endémicas y nativas de las islas Galápagos. http://www.galapagos.gob.ec/wpcontent/uploads/downloads/2016/08/Proyecto_control_y_errad.pd (accessed 29 May 2020) (In Spanish).

Earl, D.A., von Holdt, B.M., 2012. STRUCTURE HARVESTER: a website and program for visualizing STRUCTURE output and implementing the Evanno method. Conservation Genet Resour 4, 359–361. https://doi.org/10.1007/s12686-011-9548-7

Ellstrand, N.C., Elam, D.R., 1993. Population Genetic Consequences of Small Population Size: Implications for Plant Conservation. Annual Review of Ecology and Systematics 24, 217–242.

Ellstrand, N.C., Rieseberg, L.H., 2016. When gene flow really matters: gene flow in applied evolutionary biology. Evol Appl 9, 833–836. https://doi.org/10.1111/eva.12402

Emerson, B.C., 2002. Evolution on oceanic islands: molecular phylogenetic approaches to understanding pattern and process. Mol. Ecol. 11, 951–966. https://doi.org/10.1046/j.1365-294x.2002.01507.x

Evanno, G., Regnaut, S., Goudet, J., 2005. Detecting the number of clusters of individuals using the software structure: a simulation study. Molecular Ecology 14, 2611–2620. https://doi.org/10.1111/j.1365-294X.2005.02553.x

Fernández-Mazuecos, M., Jiménez-Mejías, P., Rotllan-Puig, X., Vargas, P., 2014. Narrow endemics to Mediterranean islands: Moderate genetic diversity but narrow climatic niche of the ancient, critically endangered *Naufraga* (Apiaceae). Perspectives in Plant Ecology, Evolution and Systematics 16, 190–202. https://doi.org/10.1016/j.ppees.2014.05.003

Frankham, R., 1995. Inbreeding and Extinction: A Threshold Effect. Conservation Biology 9, 792–799. https://doi.org/10.1046/j.1523-1739.1995.09040792.x

Frankham, R., 1997. Do island populations have less genetic variation than mainland populations? Heredity 78, 311–327. https://doi.org/10.1038/hdy.1997.46

Frankham, R., 1998. Inbreeding and Extinction: Island Populations. Conservation Biology 12, 665–675.

Frankham, R., Ballou, J.D., Briscoe, D.A., 2010. Introduction to Conservation Genetics. Cambridge University Press, New York.

Fridley, J.D., Grime, J.P., Bilton, M., 2007. Genetic Identity of Interspecific Neighbours Mediates Plant Responses to Competition and Environmental Variation in a Species-Rich Grassland. Journal of Ecology 95, 908–915.

Geist, D.J., Snell, Howard, Snell, Heidi, Goddard, C., Kurz, M.D., 2014. A Paleogeographic Model of the Galápagos Islands and Biogeographical and Evolutionary Implications, in: The Galápagos. American Geophysical Union (AGU), pp. 145–166. https://doi.org/10.1002/9781118852538.ch8

Gentile, G., Fabiani, A., Marquez, C., Snell, H.L., Snell, H.M., Tapia, W., Sbordon, V., 2009. An overlooked pink species of land iguana in the Galápagos. Proc. Nat. Acad. Sci. 106, 507–511.

Gentry, A.H., 1982. Patterns of Neotropical Plant Species Diversity, in: Hecht, M.K., Wallace, B., Prance, G.T. (Eds.), Evolutionary Biology: Volume 15. Springer US, Boston, MA, pp. 1–84. https://doi.org/10.1007/978-1-4615-6968-8_1

Gerlach, J., Muir, C., Richmond, M.D., 2006. The first substantiated case of trans-oceanic tortoise dispersal. Journal of Natural History 40, 2403–2408. https://doi.org/10.1080/00222930601058290

Gillespie, R., Clague, D., (Eds.), 2009. Encyclopedia of Islands. University of California Press. Retrieved June, 2020 from: www.jstor.org/stable/10.1525/j.ctt1pn90r

Gitzendanner, M.A., Weekley, C.W., Germain-Aubrey, C.C., Soltis, D.E., Soltis, P.S., 2012. Microsatellite evidence for high clonality and limited genetic diversity in *Ziziphus celata* (Rhamnaceae), an endangered, self-incompatible shrub endemic to the Lake Wales Ridge, Florida, USA. Conserv Genet 13, 223–234. https://doi.org/10.1007/s10592-011-0287-9

Grant, P.R., Grant, B.R., 2008. How and why Species Multiply: The Radiation of Darwin’s Finches. Princeton University Press.

Grenier, C., 2007. Conservación contra natura. Las Islas Galápagos. Editorial Abya Yala. (In Spanish)

Griffiths, S., 2013. Isolating microsatellite markers in the marine sponge Cinachyrella alloclada for use in community and population genetics studies (MPhil dissertation). University of Manchester, Manchester, UK.

Guerrero, A.M., Tye, A., 2009. Darwin’s Finches as Seed Predators and Dispersers. The Wilson Journal of Ornithology 121, 752–764.

Guezennec, J., Moretti, C., Simon, J.C., 2006. Natural substances in French Polynesia: utilization strategies. IRD Editions.

Guzmán, B., Heleno, R., Nogales, M., Simbaña, W., Traveset, A., Vargas, P., 2016. Evolutionary history of the endangered shrub snapdragon *(Galvezia leucantha)* of the Galápagos Islands. Diversity and Distributions 23, 247–260. https://doi.org/10.1111/ddi.12521

Hagenblad, J., Hülskötter, J., Acharya, K.P., Brunet, J., Chabrerie, O., Cousins, S.A.O., Dar, P.A., Diekmann, M., De Frenne, P., Hermy, M., Jamoneau, A., Kolb, A., Lemke, I., Plue, J., Reshi, Z.A., Graae, B.J., 2015. Low genetic diversity despite multiple introductions of the invasive plant species *Impatiens glandulifera* in Europe. BMC Genet 16. https://doi.org/10.1186/s12863-015-0242-8

Hamrick, J.L., Godt, M.J.W., 1996. Effects of Life History Traits on Genetic Diversity in Plant Species. Philosophical Transactions: Biological Sciences 351, 1291–1298.

Heleno, R.H., Olesen, J.M., Nogales, M., Vargas, P., Traveset, A., 2013. Seed dispersal networks in the Galápagos and the consequences of alien plant invasions. Proc. Biol. Sci. 280, 20122112. https://doi.org/10.1098/rspb.2012.2112

Helsen, P., Browne, R.A., Anderson, D.J., Verdyck, P., Van Dongen, S., 2009. Galápagos’ *Opuntia* (prickly pear) cacti: extensive morphological diversity, low genetic variability. Biol J Linn Soc 96, 451–461. https://doi.org/10.1111/j.1095-8312.2008.01141.x

Honnay, O., Jacquemyn, H., 2008. A meta-analysis of the relation between mating system, growth form and genotypic diversity in clonal plant species. Evol Ecol 22, 299–312. https://doi.org/10.1007/s10682-007-9202-8

Instituto Geofísico, n/d. Cámaras Volcanes Galápagos. https://www.igepn.edu.ec/islas-galapagos/content/50-islas-galapagos (accessed 18 August 2019) (in Spanish)

Instituto Nacional de Estadísticas y Censos, 2016. Galápagos tiene 25.244 habitantes según censo 2015. https://www.ecuadorencifras.gob.ec/galapagos-tiene-25-244-habitantes-segun-censo-2015/ (accessed 12 August 2019) (in Spanish)

Island Conservation, 2016. Impact Report 2015/2016. Island Conservation, Santa Cruz, CA.

Jakobsson, M., Rosenberg, N.A., 2007. CLUMPP: a cluster matching and permutation program for dealing with label switching and multimodality in analysis of population structure. Bioinformatics 23, 1801–1806. https://doi.org/10.1093/bioinformatics/btm233

Jaramillo, P., Atkinson, R., Gentile, G., 2011. Evaluating Genetic Diversity for the Conservation of the Threatened Galapagos Endemic *Calandrinia galapagosa* (Portulacaceae). Biotropica 43, 386–392. https://doi.org/10.1111/j.1744-7429.2010.00685.x

Jaramillo, P., Guézou, A., Mauchamp, A., Tye, A., 2014. CDF checklist of Galapagos flowering plants. In: Bungartz, F., Herrera, H., Jaramillo, P., Tirado, N., Jiménez-Uzcátegui, G., Ruiz, D., Guézou, A., Ziemmeck, F. (Eds.), Charles Darwin Foundation Galapagos species checklist. Charles Darwin Foundation/Fundación Charles Darwin http://www.darwinfoundation.org/datazone/checklists/vascular-plants/magnoliophyta/

Jombart, T., Ahmed, I., 2011. adegenet 1.3-1: new tools for the analysis of genome-wide SNP data. Bioinformatics 27, 3070–3071. https://doi.org/10.1093/bioinformatics/btr521

Jump, A.S., Marchant, R., Peñuelas, J., 2009. Environmental change and the option value of genetic diversity. Trends Plant Sci. 14, 51–58. https://doi.org/10.1016/j.tplants.2008.10.002

Kamvar, Z.N., Tabima, J.F., Grünwald, N.J., 2014. Poppr: an R package for genetic analysis of populations with clonal, partially clonal, and/or sexual reproduction. PeerJ 2, e281. https://doi.org/10.7717/peerj.281

Kawasaki, L., Holst, B., Bazante, G., 2017. Psidium galapageium. In: León-Yánez, S., Valencia, R., Pitmam, N., Endara, L., Ulloa, C., Navarrete, H. (Eds). Libro Rojo de Plantas Endémicas del Ecuador. Publicaciones del Herbario QCA, Pontificia Universidad Católica del Ecuador. https://bioweb.bio/floraweb/librorojo/FichaEspecie/Psidium%20galapageium

Keenan K, McGinnity P, Cross TF, Crozier WW, Prodöhl PA, 2013. DiveRsity: an R package for the estimation and exploration of population genetics parameters and their associated errors. Methods Ecol Evol 4: 782–788

Kricher, J.C., 2006. Galápagos: A Natural History. Princeton University Press, Princeton.

Kwak, M.M., Bekker, R.M., 2006. Ecology of plant reproduction: extinction risks and restoration perspectives or rare plant species. In: Waser, N.M., Ollerton, J. Waser, N.M., Ollerton, J. (Eds.), Plant-pollinator interactions: from specialization to generalization. The University of Chicago Press, 362–386.

Landrum, L.R., 2017. The Genus *Psidium* (Myrtaceae) in the State of Bahia, Brazil. Herbarium, Natural History Collections, School of Life Sciences, Arizona State University.

Latorre, O., 1991. Manuel J. Cobos, emperador de Galápagos. Fundación Charles Darwin para las Islas Galápagos. (in Spanish).

Latorre, O., 1997. Galápagos: los primeros habitantes de algunas islas. Noticias de Galapagos 56-57, 62–66. (in Spanish).

Levin, R.A., Keyes, E.M., Miller, J.S., 2015. Evolutionary Relationships, Gynodioecy, and Polyploidy in the Galápagos Endemic *Lycium minimum* (Solanaceae). International Journal of Plant Sciences 176, 197–210. https://doi.org/10.1086/679492

Loeschcke, V., Tomiuk, J., Jain, S.K. (Eds), 1994. Conservation Genetics. Experientia Supplementum Vol. 68. Springer Basel AG. 10.1007/978-3-0348-8510-2

Lombaert, E., Guillemaud, T., Thomas, C.E., Lawson Handley, L.J., Li, J., Wang, S., Pang, H., Goryacheva, I., Zakharov, I.A., Jousselin, E., Poland, R.L., Migeon, A., Van Lenteren, J., DE Clercq, P., Berkvens, N., Jones, W., Estoup, A., 2011. Inferring the origin of populations introduced from a genetically structured native range by approximate Bayesian computation: case study of the invasive ladybird *Harmonia axyridis*. Mol. Ecol. 20, 4654–4670. https://doi.org/10.1111/j.1365-294X.2011.05322.x

López-Caamal, A., Cano-Santana, Z., Jiménez-Ramírez, J., Ramírez-Rodríguez, R., Tovar-Sánchez, E., 2014. Is the insular endemic *Psidium socorrense* (Myrtaceae) at risk of extinction through hybridization? Plant Syst Evol 300, 1959–1972. https://doi.org/10.1007/s00606-014-1025-9

Lundh, J.P., 2004. Galápagos: A Brief History. http://www.galapagos.to/TEXTS/LUNDH1-1.php (accessed 20 September 2019).

MacArthur, R.H., Wilson, E.O., 1967. The Theory of Island Biogeography. Princeton University Press, Princeton.

Machado-Marques, A.M., Tuler, A.C., Carvalho, C.R., Carrijo, T.T., Ferreira, M.F. da S., Clarindo, W.R., 2016. Refinement of the karyological aspects of *Psidium guineense* (Swartz, 1788): a comparison with *Psidium guajava* (Linnaeus, 1753). Comp Cytogenet 10, 117–128. https://doi.org/10.3897/CompCytogen.v10i1.6462

Madlung, A., 2013. Polyploidy and its effect on evolutionary success: old questions revisited with new tools. Heredity (Edinb) 110, 99–104. https://doi.org/10.1038/hdy.2012.79

Mandel, J.R., Major, C.K., Bayer, R.J., Moore, J.E., 2019. Clonal diversity and spatial genetic structure in the long-lived herp, *Prairie trillium*. PLOS ONE 14(10): e0224123. https://doi.org/10.1371/journal.pone.0224123

Mayr, E., 1954. Change of genetic environment and evolution. In: Huxley, J., Hardy, A.C., Ford, E.B. (Eds.), Evolution as a Process. Allen and Unwin, 157–180.

McMullen, C.K., 1999. Flowering Plants of the Galápagos. Cornell University Press, Ithaca.

McMullen, C.K., 1990. Reproductive biology of Galapagos island angiosperms. In: Lawesson, J. E., Hamann, O., Rogers, R., Reck, G., Ochoa, H. (Eds.), Botanical research and management in Galapagos. Missouri Botanical Garden, 35–45.

Meirmans, P.G., van Tienderen, P.H.V., 2004. Genotype and Genodive: two programs for the analysis of genetic diversity of asexual organisms. Molecular Ecology Notes 4, 792–794. https://doi.org/10.1111/j.1471-8286.2004.00770.x

Meirmans, P.G., Liu, S., van Tienderen, P.H., 2018. The Analysis of Polyploid Genetic Data. J. Hered. 109, 283–296. https://doi.org/10.1093/jhered/esy006

Merlen, G., 2014. Plate tectonics, evolution, and the survival of species. In: Harpp, K. S., Mittelstaedt, E., d’Ozouville, N., Graham D. W. Harpp, K. S., Mittelstaedt, E., d’Ozouville, N., Graham D. W. (Eds.), The Galapagos: A Natural Laboratory for the Earth Sciences. John Wiley & Sons, Inc., 119–144.

Moritz, C., 2002. Strategies to protect biological diversity and the evolutionary processes that sustain it. Syst. Biol. 51, 238–254. https://doi.org/10.1080/10635150252899752

Nielsen, L.R., 2004. Molecular differentiation within and among island populations of the endemic plant *Scalesia affinis* (Asteraceae) from the Galápagos Islands. Heredity 93, 434–442. https://doi.org/10.1038/sj.hdy.6800520

Pailles, Y., Ho, S., Pires, I.S., Tester, M., Negrão, S., Schmöckel, S.M., 2017. Genetic Diversity and Population Structure of Two Tomato Species from the Galapagos Islands. Front Plant Sci 8. https://doi.org/10.3389/fpls.2017.00138

Parque Nacional Galapagos, 2016. Un sector en necesidad de renovación. http://www.carlospi.com/galapagospark/desarrollo_sustentable_agropecuario.html (accessed 31 August 2019) (in Spanish).

Peakall, R., Smouse, P.E., 2012. GenAlEx 6.5: genetic analysis in Excel. Population genetic software for teaching and research—an update. Bioinformatics 28, 2537–2539. https://doi.org/10.1093/bioinformatics/bts460

Petren, K., Grant, P.R., Grant, B.R., Keller, L.F., 2005. Comparative landscape genetics and the adaptive radiation of Darwin’s finches: the role of peripheral isolation. Mol. Ecol. 14, 2943–2957. https://doi.org/10.1111/j.1365-294X.2005.02632.x

Pluess, A.R., Stöcklin, J., 2004. Population genetic diversity of the clonal plant *Geum reptans* (Rosaceae) in the Swiss Alps. Am. J. Bot. 91, 2013–2021. https://doi.org/10.3732/ajb.91.12.2013

Porter, D.M., 1968. *Psidium* (Myrtaceae) in the Galapagos Islands. Annals of the Missouri Botanical Garden 55, 368–371. https://doi.org/10.2307/2395130

Porter, D.M., 1976. Geography and dispersal of Galapagos Islands vascular plants. Nature 264, 745–746. https://doi.org/10.1038/264745a0

Pritchard, J.K., Stephens, M., Donnelly, P., 2000. Inference of population structure using multilocus genotype data. Genetics 155, 945–959.

R Core Development Team, 2015. R: A language and environment for statistical computing. http://www.R-project.org (accessed 14 February 2020).

Reusch, T.B.H., Ehlers, A., Hämmerli, A., Worm, B., 2005. Ecosystem recovery after climatic extremes enhanced by genotypic diversity. PNAS 102, 2826–2831. https://doi.org/10.1073/pnas.0500008102

Rick, C. M., 1983. Genetic variation and evolution of Galapagos tomatoes. In: Bowman R. I., Berson, M., Leviton, A. (Eds.), Patterns of Evolution in Galapagos Organism. American Association for the Advancement of Science, 97–106.

Rick, C.M., Fobes, J.F., 1975. Allozymes of Galapagos Tomatoes: Polymorphism, Geographic Distribution, and Affinities. Evolution 29, 443–457. https://doi.org/10.1111/j.1558-5646.1975.tb00834.x

Rivas-Torres, G.F., Benítez, F.L., Rueda, D., Sevilla, C., Mena, C.F., 2018. A methodology for mapping native and invasive vegetation coverage in archipelagos: An example from the Galápagos Islands. Progress in Physical Geography: Earth and Environment. https://doi.org/10.1177/0309133317752278

Rogstad, S.H., Keane, B., Beresh, J., 2002. Genetic variation across VNTR loci in central North American *Taraxacum* surveyed at different spatial scales. Plant Ecology 161, 111–121. https://doi.org/10.1023/A:1020301011283

Rosenberg, N.A., 2004. distruct: a program for the graphical display of population structure. Molecular Ecology Notes 4, 137–138. https://doi.org/10.1046/j.1471-8286.2003.00566.x

Rumeu, B., Vargas, P., Riina, R., 2016. Incipient radiation versus multiple origins of the Galápagos *Croton scouleri* (Euphorbiaceae). Journal of Biogeography 43, 1717–1727. https://doi.org/10.1111/jbi.12753

Saghai-Maroof, M.A., Soliman, K.M., Jorgensen, R.A., Allard, R.W., 1984. Ribosomal DNA spacer-length polymorphisms in barley: mendelian inheritance, chromosomal location, and population dynamics. Proc. Natl. Acad. Sci. U.S.A. 81, 8014–8018. https://doi.org/10.1073/pnas.81.24.8014

Sakai, A.K., Allendorf, F.W., Holt, J.S., Lodge, D.M., Molofsky, J., With, K.A., Baughman, S., Cabin, R.J., Cohen, J.E., Ellstrand, N.C., McCauley, D.E., O’Neil, P., Parker, I.M., Thompson, J.N., Weller, S.G., 2001. The Population Biology of Invasive Species. Annu. Rev. Ecol. Syst. 32, 305–332. https://doi.org/10.1146/annurev.ecolsys.32.081501.114037

Sakai, A.K., Wagner, W.L., Ferguson, D.M., Herbst, D.R., 1995. Origins of Dioecy in the Hawaiian Flora. Ecology 76, 2517–2529. https://doi.org/10.2307/2265825

Schmitz, P., Cibois, A., Landry, B., 2007. Molecular phylogeny and dating of an insular endemic moth radiation inferred from mitochondrial and nuclear genes: the genus *Galagete* (Lepidoptera: Autostichidae) of the Galapagos Islands. Mol. Phylogenet. Evol. 45, 180–192. https://doi.org/10.1016/j.ympev.2007.05.010

Scobie, A.R., Wilcock, C.C., 2009. Limited mate availability decreases reproductive success of fragmented populations of Linnaea borealis, a rare, clonal self-incompatible plant. Ann Bot 103, 835–846. https://doi.org/10.1093/aob/mcp007

Sequeira, A.S., Sijapati, M., Lanteri, A.A., Roque Albelo, L., 2008. Nuclear and mitochondrial sequences confirm complex colonization patterns and clear species boundaries for flightless weevils in the Galápagos archipelago. Philosophical Transactions of the Royal Society B: Biological Sciences 363, 3439–3451. https://doi.org/10.1098/rstb.2008.0109

Shaw, K.L., Gillespie, R.G., 2016. Comparative phylogeography of oceanic archipelagos: Hotspots for inferences of evolutionary process. Proc Natl Acad Sci U S A 113, 7986–7993. https://doi.org/10.1073/pnas.1601078113

Shirk, R.Y., Hamrick, J.L., Zhang, C., Qiang, S., 2014. Patterns of genetic diversity reveal multiple introductions and recurrent founder effects during range expansion in invasive populations of *Geranium carolinianum* (Geraniaceae). Heredity (Edinb) 112, 497–507. https://doi.org/10.1038/hdy.2013.132

Silvertown, J., 2008. The Evolutionary Maintenance of Sexual Reproduction: Evidence from the Ecological Distribution of Asexual Reproduction in Clonal Plants. International Journal of Plant Sciences 169, 157–168. https://doi.org/10.1086/523357

Singhal, V.K., Gill, B.S., Bir, S.S., 1985. Cytology of woody species. Proc. Indian Acad. Sci. (Plant Sci.) 94, 607–618. https://doi.org/10.1007/BF03053228

Sitther, V., Zhang, D., Harris, D.L., Yadav, A.K., Zee, F.T., Meinhardt, L.W., Dhekney, S.A., 2014. Genetic characterization of guava *(Psidium guajava* L.) germplasm in the United States using microsatellite markers. Genet Resour Crop Evol 61, 829–839. https://doi.org/10.1007/s10722-014-0078-5

Smith, J.M.B., 2009. Dispersal of plants and animals to oceanic islands. In: Wolanski, E. (Ed.), Oceans and aquatic ecosystems Volume II. EOLSS, 269–283.

Soltis, P.S., Soltis, D.E., 2000. The role of genetic and genomic attributes in the success of polyploids. Proceedings of the National Academy of Sciences of the United States of America. https://doi.org/10.1073/pnas.97.13.7051

Stuessy, T.F., Jakubowsky, G., Gómez, R.S., Pfosser, M., Schlüter, P.M., Fer, T., Sun, B.-Y., Kato, H., 2006. Anagenetic evolution in island plants. Journal of Biogeography 33, 1259–1265. https://doi.org/10.1111/j.1365-2699.2006.01504.x

Stuessy, T.F., Takayama, K., López-Sepúlveda, P., Crawford, D.J., 2014. Interpretation of patterns of genetic variation in endemic plant species of oceanic islands. Bot J Linn Soc 174, 276–288. https://doi.org/10.1111/boj.12088

Su, Y., Wang, T., Deng, F., 2010. Contrasting genetic variation and differentiation on Hainan Island and the Chinese mainland populations of *Dacrycarpus imbricatus* (Podocarpaceae). Biochemical Systematics and Ecology 38, 576–584. https://doi.org/10.1016/j.bse.2010.07.003

Sundqvist, L., Keenan, K., Zackrisson, M., Prodöhl, P., Kleinhans, D., 2016. Directional genetic differentiation and relative migration. Ecology and Evolution 6, 3461–3475. https://doi.org/10.1002/ece3.2096

Takayama, K., López Sepúlveda, P., Kohl, G., Novak, J., Stuessy, T.F., 2013. Development of microsatellite markers in *Robinsonia* (Asteraceae) an endemic genus of the Juan Fernández Archipelago, Chile. Conservation Genet Resour 5, 63–67. https://doi.org/10.1007/s12686-012-9734-2

Takayama, K., López-Sepúlveda, P., Greimler, J., Crawford, D.J., Peñailillo, P., Baeza, M., Ruiz, E., Kohl, G., Tremetsberger, K., Gatica, A., Letelier, L., Novoa, P., Novak, J., Stuessy, T.F., 2015. Genetic consequences of cladogenetic vs. anagenetic speciation in endemic plants of oceanic islands. AoB Plants 7. https://doi.org/10.1093/aobpla/plv102

Torres, M. de L., Gutiérrez, B., 2018. A Preliminary Assessment of the Genetic Diversity and Population Structure of Guava, *Psidium guajava,* in San Cristobal. In: Torres, M. de L., Mena, C.F. (Eds.), Understanding Invasive Species in the Galapagos Islands: From the Molecular to the Landscape, Social and Ecological Interactions in the Galapagos Islands. Springer International Publishing, Cham, pp. 3–17. https://doi.org/10.1007/978-3-319-67177-2_1

Traveset, A., Heleno, R., Chamorro, S., Vargas, P., McMullen, C.K., Castro-Urgal, R., Nogales, M., Herrera, H.W., Olesen, J.M., 2013. Invaders of pollination networks in the Galápagos Islands: emergence of novel communities. Proceedings of the Royal Society B: Biological Sciences 280, 20123040. https://doi.org/10.1098/rspb.2012.3040

Tuler, A.C., Carrijo, T.T., Peixoto, A.L., Garbin, M.L., da Silva Ferreira, M.F., Carvalho, C.R., Spadeto, M.S., Clarindo, W.R., 2019. Diversification and geographical distribution of *Psidium* (Myrtaceae) species with distinct ploidy levels. Trees 33, 1101–1110. https://doi.org/10.1007/s00468-019-01845-2

Tye, A., Atkinson, R., Carrión, V., 2007. Increase in the number of introduced plant species in Galapagos (Galapagos Report 2006-2007). CDF, GNP and INGALA, Puerto Ayora, Galápagos, Ecuador.

Urquía, D., Gutierrez, B., Pozo, G., Pozo, M.J., Espín, A., Torres, M. de L., 2019. *Psidium guajava* in the Galapagos Islands: Population genetics and history of an invasive species. PLOS ONE 14, e0203737. https://doi.org/10.1371/journal.pone.0203737

Valdebenito, H., 2018. A Study Contrasting Two Congener Plant Species: *Psidium guajava* (introduced guava) and *P. galapageium* (Galapagos guava) in the Galapagos Islands. In: Torres, M. de L., Mena, C.F. (Eds.), Understanding Invasive Species in the Galapagos Islands: From the Molecular to the Landscape, Social and Ecological Interactions in the Galapagos Islands. Springer International Publishing, Cham, pp. 47–68. https://doi.org/10.1007/978-3-319-67177-2_1

Vellend, M., Geber, M.A., 2005. Connections between species diversity and genetic diversity. Ecology Letters 8, 767–781. https://doi.org/10.1111/j.1461-0248.2005.00775.x

Villagómez, D.R., Toomey, D.R., Hooft, E.E.E., Solomon, S.C., 2007. Upper mantle structure beneath the Galápagos Archipelago from surface wave tomography. Journal of Geophysical Research: Solid Earth 112. https://doi.org/10.1029/2006JB004672

Wallis, G.P., Trewick, S.A., 2009. New Zealand phylogeography: evolution on a small continent. Mol. Ecol. 18, 3548–3580. https://doi.org/10.1111/j.1365-294X.2009.04294.x

Ward, S.M., Gaskin, J.F., Wilson, L.M., 2008. Ecological Genetics of Plant Invasion: What Do We Know? Ipsm 1, 98–109. https://doi.org/10.1614/IPSM-07-022.1

Ward, R.G., Brookfield, M., 1992. Special Paper: The Dispersal of the Coconut: Did It Float or Was It Carried to Panama? Journal of Biogeography 19, 467–480. https://doi.org/10.2307/2845766

Wendel, J.F., Percival, A.E., 1990. Molecular divergence in the Galapagos Islands— Baja California species pair, *Gossypium klotzschianum* and *G. davidsonii* (Malvaceae). Pl Syst Evol 171, 99–115. https://doi.org/10.1007/BF00940598

Wendel, J.F., Percy, R.G., 1990. Allozyme diversity and introgression in the Galapagos Islands endemic *Gossypium darwinii* and its relationship to continental *G. barbadense*. Biochemical Systematics and Ecology 18, 517–528. https://doi.org/10.1016/0305-1978(90)90123-W

Wendel, J.F., Cronn, R.C., 2003. Polyploidy and the evolutionary history of cotton. Advances in Agronomy. 87: 139–186.

Whittaker, R. J., 1998. The human impact on islands ecosystems – the lighthouse keeper’s cat and other stories. In: Whittaker, R. J., Fernández Palacios, J. M. (Eds.), Island Biography: Ecology, Evolution and Conservation. Oxford University Press, 237–265.

Whittaker, R.J., Fernandez-Palacios, J.M., 2007. Island Biogeography: Ecology, Evolution, and Conservation. OUP Oxford.

Wickham, H., 2009. ggplot2: Elegant Graphics for Data Analysis, Springer-Verlag, New York. https://doi.org/10.1007/978-0-387-98141-3

Wiggins, I.L., Porter, D.M., Anderson, E.F., 1971. Flora of the Galapagos Islands. Stanford University Press.

Witter, M.S., Carr, G.D., 1988. Adaptive radiation and genetic differentiation in the Hawaiian silversword alliance (Compositae: Madiinae). Evolution 42, 1278–1287. https://doi.org/10.1111/j.1558-5646.1988.tb04187.x

Wright, S., 1951. The Genetical Structure of Populations. Annals of Eugenics 15, 323–354. https://doi.org/10.1111/j.1469-1809.1949.tb02451.x

