## Appendix_A for "Understanding the genetic diversity of the guayabillo (*Psidium galapageium*), an endemic plant of the Galapagos Islands"

Table A1. Sampling sites with its coordinates and altitude, and number of individuals collected there.

| Island | | Sampling Site | Coordinates | | Altitude (masl) | Number of individuals | Total |
|---|---|---|---|---|---|---|---|
| Isabela | 1 | Ricardo García | 0° 51.308'S | 91° 00.023'W | 148 | 15 | 86 |
| | 2 | San Joaquín | 0° 49.130'S | 91° 01.304'W | 379 | 18 | |
| | 3 | El Basurero | 0° 52.359'S | 91° 00.137'W | 125 | 8 | |
| | 4 | Finca Morocho | 0° 51.040'S | 90° 59.442'W | 139 | 20 | |
| | 5 | El Mango | 0° 53.135'S | 91° 00.430'W | 127 | 6 | |
| | 6 | Cerro Grande | 0° 49.506'S | 91° 00.215'W | 258 | 19 | |
| Santa Cruz | 1 | Granillo Rojo | 0° 36.931'S | 90° 22.048'W | 574 | 14 | 87 |
| | 2 | Salasaca | 0° 37.916'S | 90° 26.188'W | 382 | 5 | |
| | 3 | Camote | 0° 38.279'S | 90° 17.448'W | 442 | 9 | |
| | 4 | Garrapatero | 0° 40.367'S | 90° 14.460'W | 132 | 12 | |
| | 5 | Bellavista | 0° 41.558'S | 90° 19.037'W | 164 | 34 | |
| | 6 | El Chato | 0° 41.907'S | 90° 24.118'W | 228 | 13 | |
| San Cristobal | 1 | Galapaguera | 0° 54.893'S | 89° 26.106'W | 109 | 5 | 35 |
| | 2 | Camino a Opuntias | 0° 56.120'S | 89° 32.819'W | 124 | 5 | |
| | 3 | Perimetral | 0° 55.917'S | 89° 32.923'W | 150 | 4 | |
| | 4 | Cerro Verde | 0° 54.416'S | 89° 26.513'W | 206 | 5 | |
| | 5 | Las Goteras | 0° 53.058'S | 89° 26.135'W | 311 | 5 | |
| | 6 | Cerro Gato | 0° 55.452'S | 89° 28.172'W | 161 | 5 | |
| | 7 | Centro de Reciclaje | 0° 54.724'S | 89° 34.794'W | 138 | 6 | |
| | Total | | | | | | 208 |

Table A2. Null allele frequencies (using both, a 0.5 and a 0.65 rate of selfing) and PICs for the two isoloci of each analyzed SSR locus.

| | **Null allele freq. (SELFING RATE=0.5)** | | **Null allele freq. (SELFING RATE=0.65)** | | **PIC** | |
|---|---|---|---|---|---|---|
| **Locus** | **Isolocus 1** | **Isolocus 2** | **Isolocus 1** | **Isolocus 2** | **Isolocus 1** | **Isolocus 2** |
| **GYB3** | 0.313 | 0.317 | 0.312 | 0.312 | 0.751 | 0.718 |
| **GYB4** | 0.225 | 0.175 | 0.229 | 0.174 | 0.411 | 0.545 |
| **GYB5** | 0.134 | 0.301** | 0.146 | 0.281** | 0.006 | 0.434** |
| **GYB7** | 0.263 | 0.261 | 0.257 | 0.253 | 0.494 | 0.466 |
| **GYB8** | 0.286 | 0.197 | 0.271 | 0.194 | 0.726 | 0.623 |
| **GYB9** | 0.070 | 0.180 | 0.071 | 0.171 | 0.674 | 0.753 |
| **GYB14** | 0.337* | 0.347* | 0.325* | 0.334* | 0.646* | 0.653* |
| **GYB18** | 0.388* | 0.372* | 0.383* | 0.367* | 0.678* | 0.749* |
| **GYB21** | 0.279 | 0.306 | 0.264 | 0.293 | 0.708 | 0.620 |
| **GYB22** | 0.224 | 0.194 | 0.218 | 0.196 | 0.358 | 0.502 |
| **GYB23** | 0.183 | 0.234 | 0.187 | 0.227 | 0.150 | 0.613 |
| **GYB27** | 0.192 | 0.346* | 0.189 | 0.333* | 0.651 | 0.500* |
| **GYB29** | 0.173 | 0.314 | 0.175 | 0.301 | 0.384 | 0.701 |

*Null allele frequency >>0.3 for both selfing rates, discarded from further analyses
**Discarded due to monomorphism.

Table A3. Genetic diversity information of the analyzed *Psidium galapageium* populations from Isabela, Santa Cruz and San Cristobal islands, after systematic downsampling in the Isabela and Santa Cruz samples: Number of individuals genotyped from each island (N), number of alleles found (A), number of private alleles (PA), mean allelic richness after rarefaction (AR), observed heterozygosity ($H_O$), expected heterozygosity/gene diversity ($H_E$) and $F_{ST}$ global value for each island population. Overall results along the three islands are also shown.

| **Island** | **N** | **A*** | **PA*** | **$H_O$ [a]** | **$H_E$ [a]** |
|---|---|---|---|---|---|
| Isabela | 35 | 118 (97) | 52 (38) | 0.122 | 0.588 |
| Santa Cruz | 35 | 84 (59) | 17 (9) | 0.156 | 0.412 |
| San Cristobal | 35 | 70 (60) | 12 (5) | 0.119 | 0.283 |
| **Overall** | 105 | 161 | - | 0.141 | 0.465 |

* Values between brackets are the number of alleles or private alleles with a frequency >0.05 within the corresponding island population.
[a]indicates average across the 15 SSRs analyzed.
[s]standardized for N=35

Table A4. Pairwise and global $F_{ST}$ values between the *Psidium galapageium* populations from the three islands, after systematic downsampling in the Isabela and Santa Cruz samples.

| | **Isabela** | **Santa Cruz** |
|---|---|---|
| **Santa Cruz** | 0.209 | - |
| **San Cristobal** | 0.212 | 0.328 |
| **Global** | 0.319 | |

Table A5. Pairwise and global $F_{ST}$ values between the *Psidium galapageium* clusters defined from the STRUCTURE software and PCoA groupings.

| | **Isabela** | **Santa Cruz 1** | **Santa Cruz 2** |
|---|---|---|---|
| **Santa Cruz 1** | 0.228 | - | |
| **Santa Cruz 2** | 0.132 | 0.166 | - |
| **San Cristobal** | 0.180 | 0.290 | 0.222 |
| **Global** | 0.314 | | |

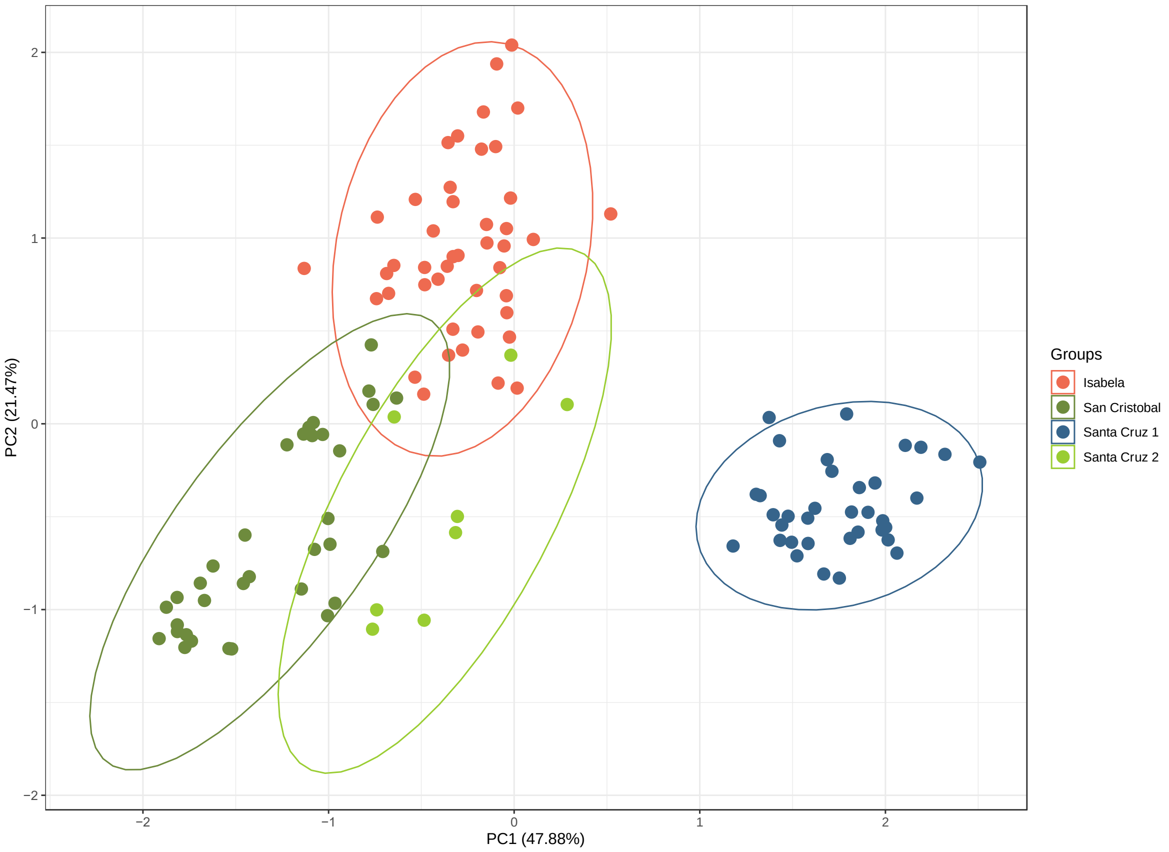


Fig. A1. PCoA based on the Lynch distances (after systematic downsampling in the Isabela and Santa Cruz samples) found between the *Psidium galapageium* individuals sampled in the three islands: Isabela, San Cristobal and Santa Cruz. For Santa Cruz, both genetic clusters are indicated (Santa Cruz 1 and Santa Cruz 2).

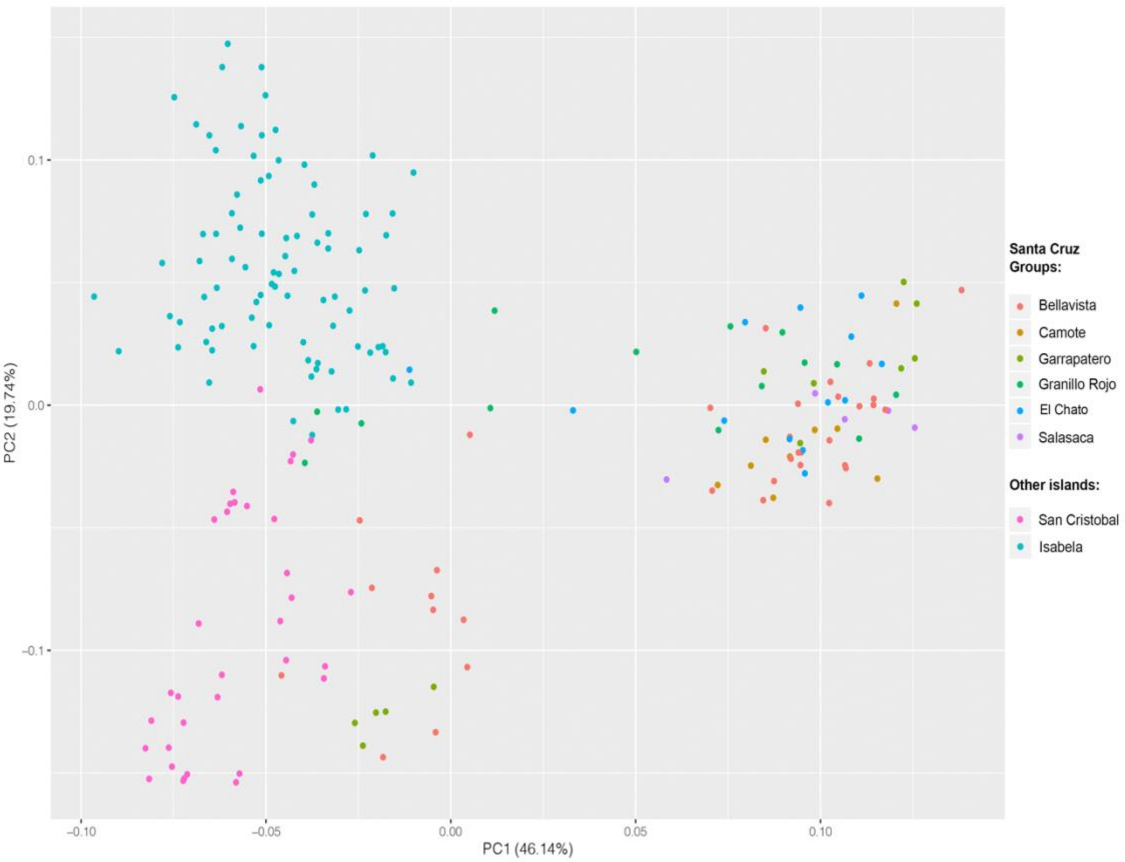


Fig. A2. PCoA based on the Lynch distances found between the *Psidium galapageium* individuals sampled in the three islands. Here, the different subpopulations of Santa Cruz are represented in different colors to show how some individuals (from Granillo Rojo, Garrapatero and Bellavista locations) are grouped with the individuals from Isabela and San Cristobal rather than with the other individuals from Santa Cruz.

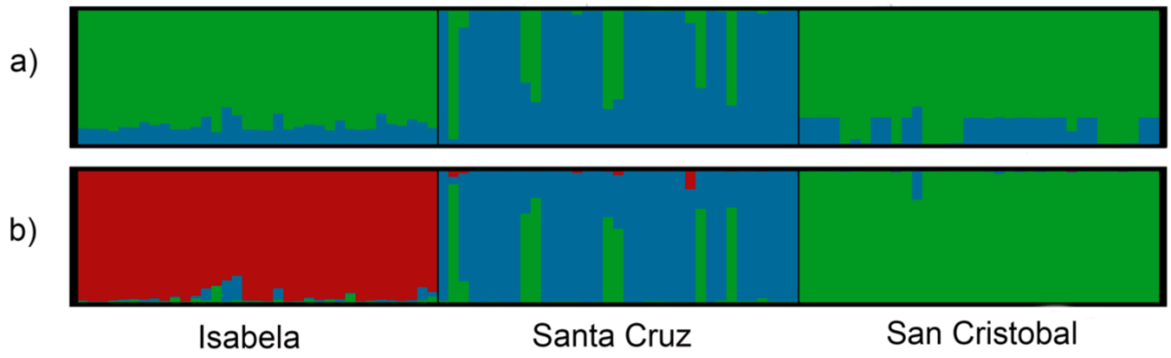


Fig. A3. Results of the Bayesian analysis of population structure (Software STRUCTURE) under the Admixture model, after systematic downsampling in the Isabela and Santa Cruz samples. The results are indicated for a) K=2, and b) K = 3 which is the optimum K value ($\Delta K = 250.69$). These values of K correspond to the clusters or lineages (represented by different colors) in which are grouped the *Psidium galapageium* individuals sampled in Isabela, Santa Cruz and San Cristobal islands.

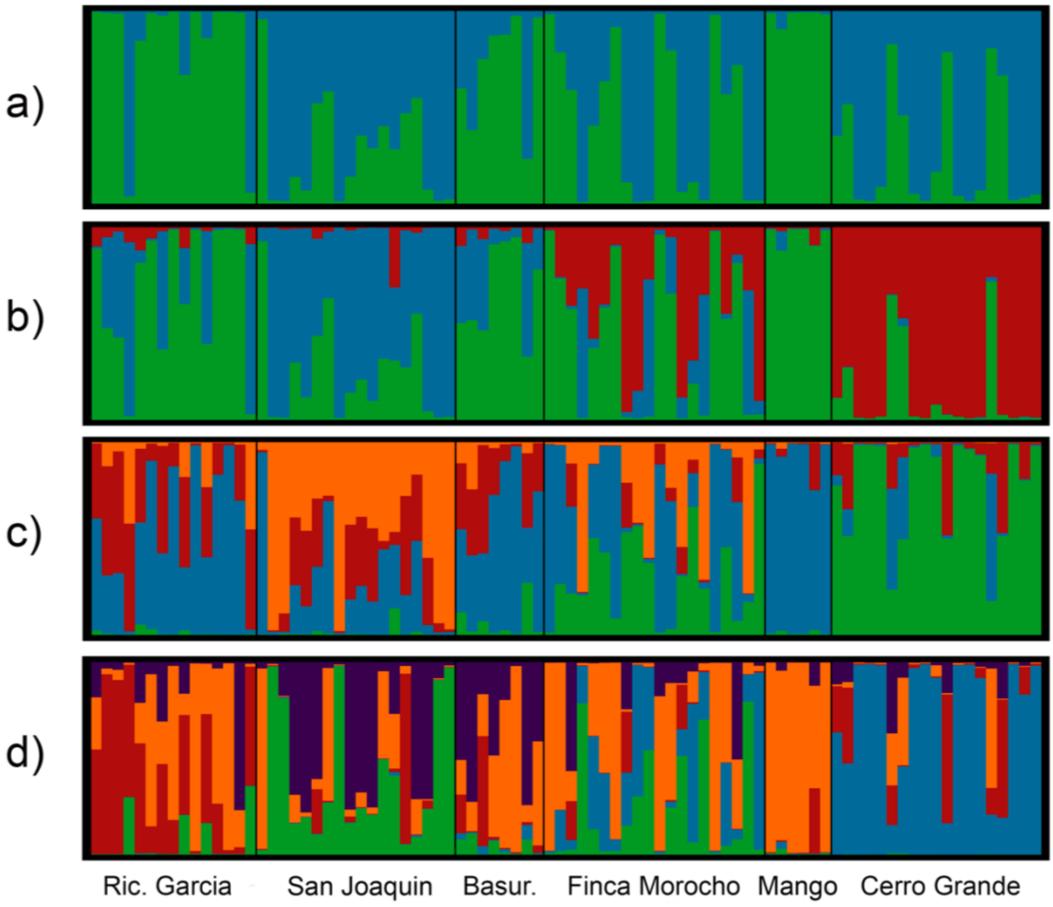


Fig. A4. Population structure Bayesian analysis results, among localities in Isabela Island (Admixture model). a) K=2, b) K=3, c) K=4, d) K=5. The optimum K value in this case was K=2 (ΔK=1195.71).

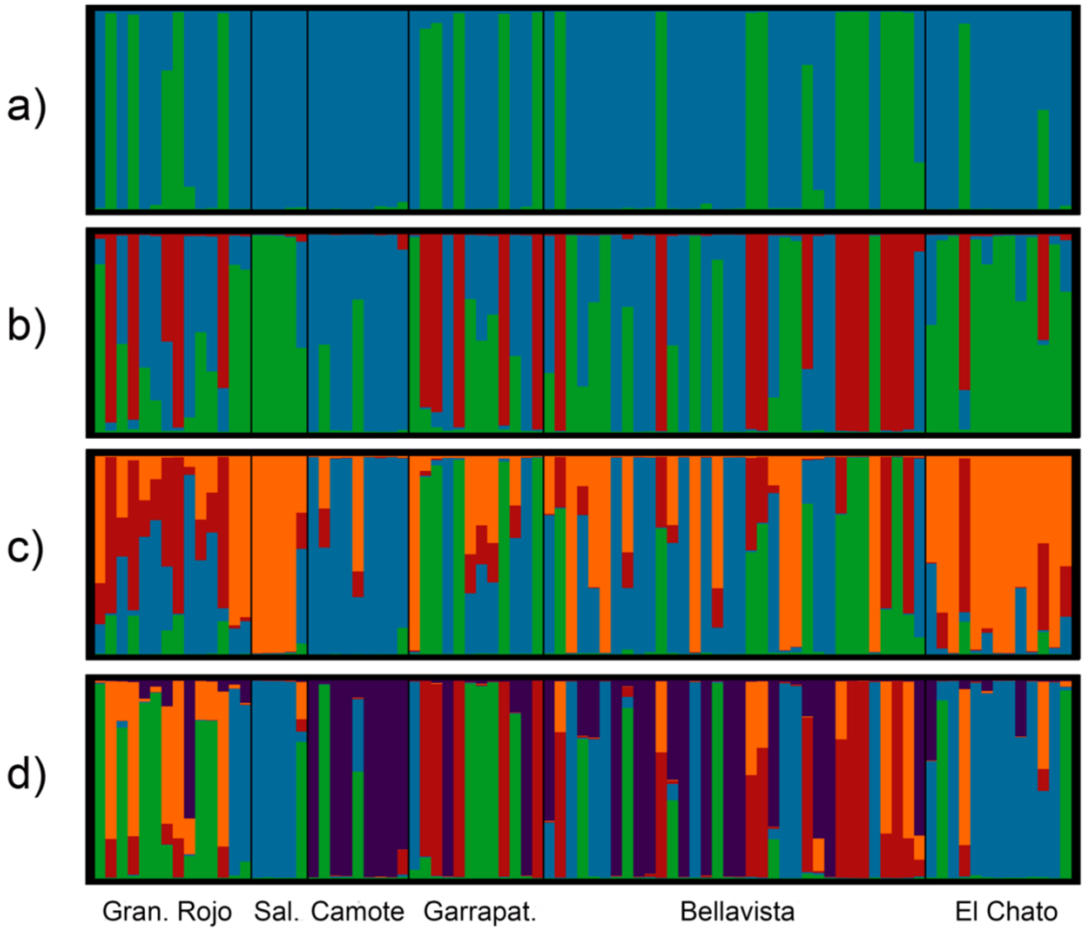


Fig. A5. Population structure Bayesian analysis results, among localities in Santa Cruz Island (Admixture model). a) K=2, b) K=3, c) K=4, d) K=5. The optimum K value in this case was K=2 (ΔK=1177.14).

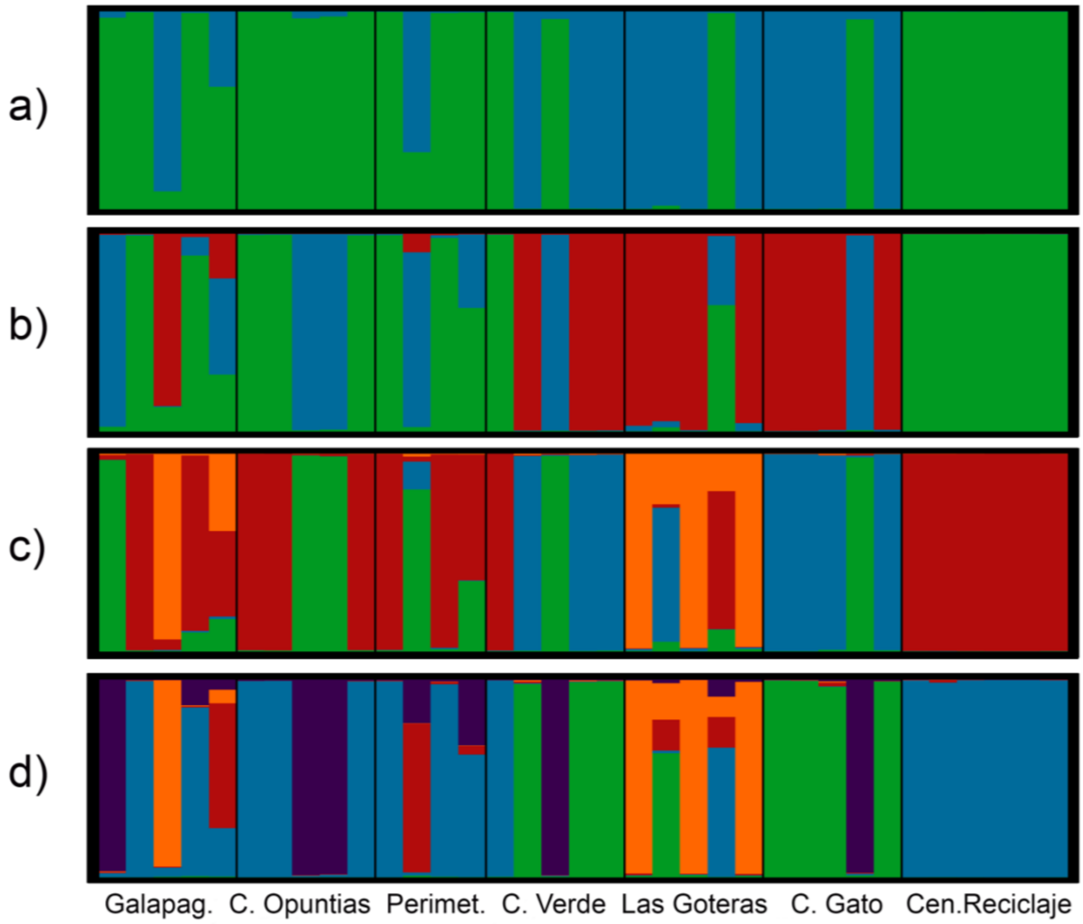


Fig. A6. Population structure Bayesian analysis results, among localities in San Cristobal Island (Admixture model). a) K=2, b) K=3, c) K=4, d) K=5. The optimum K value in this case was K=2 (ΔK= 533.70).
